# Cholesteryl Hemiazelate Induces Lysosome Dysfunction and Exocytosis in Macrophages

**DOI:** 10.1101/2021.01.05.422575

**Authors:** Neuza Domingues, Rita Diogo Almeida Calado, Patrícia H. Brito, Rune Matthiesen, José Ramalho, Maria I. L. Soares, Telmo Pereira, Luis Oliveira, José R. Vicente, Louise H. Wong, Soo Min Cho, Ines C. M. Simões, Julio Sampaio, Christian Klose, Michal A. Surma, Manuel S. Almeida, Gustavo Rodrigues, Pedro Araújo-Gonçalves, Jorge Ferreira, Kai Simons, Teresa M. V. D. Pinho e Melo, Andrew Peden, Claudia Guimas Almeida, Clare E. Futter, Anthony H. Futerman, Winchil L.C. Vaz, Otilia V. Vieira

**Author notes:** Address correspondence to: Dr. Otilia V. Vieira, CEDOC, NOVA Medical School | Faculdade de Ciências Médicas, Universidade NOVA de Lisboa, 1169-056 Lisboa, Portugal. UCIBIO-REQUIMTE, Departamento de Ciências da Vida, Faculdade de Ciências e Tecnologia, Universidade Nova de Lisboa, 2829-516 Caparica, Portugal (Patrícia H. Brito). Centre de Recherche, Institut Curie, 26 rue d’Ulm, 75248 Paris Cedex 05, France (Julio Sampaio).

## Abstract

**OBJECTIVE:** A key event in atherogenesis is the formation of lipid-loaded macrophages, lipidotic cells, which exhibit irreversible accumulation of undigested modified low-density lipoproteins in lysosomes. This event culminates with the loss of cell homeostasis, inflammation and cell death. In this study we propose to identify the chemical etiological factors and understanding the molecular and cellular mechanisms responsible for the impairment of lysosome function in macrophages.

**APPROACH AND RESULTS:** Using shotgun lipidomics we have discovered that a family of oxidized lipids (cholesteryl hemiesters, ChE), end products of oxidation of polyunsaturated cholesteryl esters, occurs at higher concentrations in the plasma of two cohorts of cardiovascular disease patients than in the plasma of a control cohort. Macrophages exposed to the most prevalent ChE, cholesteryl hemiazelate (ChA) exhibit lysosome enlargement, peripheral lysosomal positioning, lysosome dysfunction and lipidosis which are irreversible. The transcriptomic profile of macrophages exposed to ChA indicates that the lysosome pathway is deeply affected and is well correlated with lysosome phenotypic and functional changes. Interestingly, the dysfunctional peripheral lysosomes are more prone to fuse with the plasma membrane, secreting their undigested luminal content into the extracellular milieu with potential consequences to the pathology.

**CONCLUSION:** We identify ChA not only as one of the molecules involved in the etiology of irreversible lysosome dysfunction culminating with lipidosis but also as a promoter of exocytosis of the dysfunctional lysosomes. The latter event is a new mechanism that may be important in the pathogenesis of atherosclerosis.

## INTRODUCTION

Atherosclerosis is a progressive insidious chronic disease that underlies most of the cardiovascular complications worldwide including myocardial infarction, stroke, and heart failure.^1^ The factors that influence the initiation and progression of atherosclerotic lesions are extremely complex and still relatively poorly understood ^2^. One of the early characteristics of atherogenesis, observed in lesions of different origin, is the sequestration of lipids by macrophages in the arterial *intima* ^3-9^. This results from ingestion of modified low-density lipoproteins (LDL) accumulated in the vascular wall by macrophages, and intracellular accumulation of their lipids, leading to the formation of intravascular “fatty streaks”. Although this process is initially reversible, it becomes irreversible as a result of inadequate processing of the internalized lipids by the macrophages. The result is an accumulation of undigested cargo in the macrophage lysosomal compartment, leading to disruption of both lysosome function and cellular homeostasis ^10-13^.

LDL uptake is a highly regulated system with feedback mechanisms limiting excessive uptake and lipid overload ^14, 15^. In atherogenesis, there is a consensus that foam cell formation requires entrapment of LDL within the arterial intima followed by some modification of the entrapped particles. One modification that LDL particles undergo is oxidation ^16-20^. This modified LDL is internalized through unregulated scavenger receptor pathways or through micropinocytosis ^21^. Moreover, increasing evidence suggest that macrophages loaded with oxidized LDL (Ox-LDL) have disrupted lysosomal function, which result in the accumulation of intralysosomal free cholesterol (FC), cholesteryl esters and oxidized apolipoprotein B (ApoB) ^5, 11, 12, 22^. The accumulation of undigested Ox-LDL, within the lysosomal compartment, not only partially inactivates lysosomal cathepsins but also destabilizes lysosomal pH and causes relocation of lysosomal enzymes to the cytosol contributing to inflammation and apoptosis ^10, 13, 23-29^. However, due to the complexity of the chemical composition of Ox-LDL, the exact etiological agent(s) and molecular mechanisms responsible for the impairment of lysosome function in macrophages have not been identified.

Here we identify cholesteryl hemiesters (ChE), the expected oxidation end-products of the polyunsaturated fatty acid esters of cholesterol, in the plasma of cardiovascular disease (CVD) patients and address the role of the most prevalent ChE found in the plasma of CVD patients, ChA (the cholesteryl hemiester of azelaic acid), as an inducer of lysosomal dysfunction in macrophages. Our data show that lysosomes are targeted by ChA culminating with lipid accumulation and changes in their intracellular distribution and morphology. One important aspect of this work is that these lysosomes are more prone to exocytosis leading us to postulate that as an emergency mechanism, affected macrophages secrete their lysosomal contents which in turn can re-initiate inflammation and fuel the continuation of the atherogenic pathology.

## METHODS

The authors declare that all supporting data are available within the article and in the Data Supplement. The description of the main reagent is available in Supplementary Table I.

### Plasma samples

Blood samples were obtained from CVD patients and healthy individuals after explaining the purpose of the study and obtaining written informed-consent from them or their legal representatives. The entire process was approved by the Ethical Review Board of the Faculty of Medicine of the New University of Lisbon and the Ethics Committee for Health of the Centro Hospitalar de Lisboa Ocidental, Hospital Santa Cruz. All experiments were performed in accordance with the guidelines and regulations. The details for the different cohorts used and sample preparation were already been described elsewhere ^30^. After collection, plasma samples were immediately frozen at −80 °C.

### Cholesteryl hemiesters synthesis

Cholesteryl hemiesters were prepared following a general procedure described in the literature for the synthesis of cholesteryl hemisuccinate (ChS) ^31^. However, in order to optimize reaction conditions, molar equivalents of the reactants, reaction time and purification conditions were adapted.

ChA, was synthesized from the reaction of commercially available cholesterol with freshly prepared azelaic anhydride, which was obtained by reacting azelaic acid with acetyl chloride, as described elsewhere ^32^. Cholesterol was reacted with 2.6 molar equivalents of azelaic anhydride in dry pyridine under reflux for 7 h (Supplementary Figure I). Purification by flash chromatography with chloroform/methanol/ammonia (50:5:0.25), followed by recrystallization from ethanol, gave the target ChA as a white solid in 57 % yield. Cholesteryl hemiglutarate (ChG, cholesteryl-O-(4-carboxybutanoyl)) was prepared by the reaction of commercially available cholesterol with 1.7 molar equivalents of commercially available glutaric anhydride in dry pyridine under reflux for 6 h (Supplementary Figure I). Trituration of the crude product with methanol, followed by purification by flash chromatography with chloroform/methanol/ammonia (50:5:0.25), gave the target ChG as a white solid in 37 % yield. The detailed experimental procedure and characterization data for ChA and ChG are available in the Supplementary Material.

### Lipid extraction, MS lipidomics, data acquisition and analysis

Mass spectrometry-based lipid analysis was performed at Lipotype GmbH (Dresden, Germany) as previously described ^33^. Briefly, 50 µL of diluted plasma (equivalent to 1 μL of undiluted plasma) was mixed with 130 μL of 150 mM ammonium bicarbonate solution and 810 μL of methyl tert-butyl ether/methanol (7:2, v/v) was added. 21 µL of an internal standard mixture was pre-mixed with the mixture of organic solvents. The internal standard mixture covered the major lipid classes present in plasma as described previously ^30^. Additionally, 100 pmol per sample of ChS was added for quantification of ChE. The plate was then sealed with a teflon-coated lid, shaken at 4°C for 15 min, and spun down (3000 g, 5 min) to facilitate separation of the liquid phases. One hundred microliters of the organic phase was transferred to an infusion plate and dried in a speed vacuum concentrator. Dried lipids were re-suspended in 40 μL of 7.5 mM ammonium acetate in chloroform/methanol/propanol (1:2:4, v/v/v) and the wells were sealed with an aluminum foil to avoid evaporation and contamination during infusion. All liquid handling steps were performed using a Hamilton STARlet robotic platform with the Anti Droplet-Control feature for pipetting of organic solvents. Samples were analysed by direct infusion in a QExactive mass spectrometer (Thermo Fisher Scientific) equipped with a TriVersa NanoMate ion source (Advion Biosciences). 5 µL were infused with gas pressure and voltage set to 1.25 psi and 0.95 kV, respectively. We scanned for the m/z 400–650 in FTMS − (Rm/z = 200 = 280 000) for 15 s with lock mass activated at a common background (m/z = 529.46262) to detect ChE as deprotonated anions. Automatic gain control was set to 106 and ion trap filling time was set to 50 ms. ChE were identified and quantified by their accurate intact mass from the FTMS spectra with Lipotype Xplorer, a proprietary software developed from LipidXplorer ^34, 35^. The lipidomic analysis of cells extracts was carried out as previously described ^36^.

### Liposomes preparation

POPC and ChA, cholesteryl linoleate and cholesterol, were mixed at a 35:65 molar ratio. The detailed protocol for liposomes preparation was described previously by our group ^36^.

### Cell culture

RAW 264.7 cells (ATCC) were maintained and cultivated as described ^37^. Macrophages were then incubated with POPC-, POPC-Chol-, POPC-ChA-liposomes or cholesteryl linoleate emulsions for up to 72 h at different concentrations, as indicated in the figure legends.

Cell toxicity was evaluated by the MTT test as described previously ^36^.

### MARS-Seq

After cells were exposed to the desired treatment, mRNA was isolated using TRIzol and RNeasy Mini Kit. For RNA sequencing, we used a derivation of MARS-seq for the generation of RNA-Seq libraries ^38^, developed for single-cell RNA-seq. RNA-Seq with R1 65 bases in length were sequenced using Illumina NextSeq® 500 High Output v2 Kit (75 cycles). Raw reads were aligned to the genome (NCBI37/mm10) using Hisat (version 0.1.6) and only reads with unique mapping were considered for further analysis. BBCU-NGS pipelines supplied by the Bioinformatics unit in Weizmann Institute of Science ^39^ were used for further analysis of the corrected reads.

### Differentially gene expression (DGE) analysis

RNAseq expression data were analysed using the edgeR version 3.24.3 R package ^40^. Two treatment conditions POPC and ChA with four paired replicates each, were analysed. Genes were mapped to Entrez codes using org.Mm.eg.db version 3.7.0 R package ^41^ and filtered to exclude genes with overall low coverage. Genes with less than 4 counts per million (c.p.m.) in at least 75 % of the samples of each group were filtered out. From an initial dataset of 9470 different loci our final dataset included 8385 genes. The RNA count data for each library were normalized using a trimmed mean of M-values (TMM) ^42^ between each pair of samples as applied in the calcNormFactors function of edgeR. This normalization finds a set of scaling factors for the library sizes that minimize the log-fold changes between the samples for most genes. Normalized count data were fitted to a negative binomial generalized linear model with function glmQLFit(). Differential expression between treatments adjusting for baseline differences between replicates was estimated by applying a quasi-likelihood (QL) F test using the function glmQLFTest() and glmTreat(). Genes with False Discovery Rate (FDR) < 0.05 and absolute log2FC significantly higher than 1.1 were defined as significant differentially expressed.

### Gene set enrichment analysis

Using the function kegg.gsets from the R package GAGE ^43^ 238 canonical signaling and metabolism pathways present in KEGG database for mouse ^44^ were obtained. Gene set enrichment analysis was performed using fry() function from the edgeR package. The data that support the findings of this study and the scripts used for analyses are available from the corresponding author upon reasonable request. The data that support the findings of this study and the scripts used for analyses are available from the corresponding author upon reasonable request.

### Fluorescence microscopy, stainings, image acquisition and analysis

For immunofluorescence (IF) staining, RAW cells were grown in presence of the desired lipids, at the desired concentrations and times, as indicated in the figure legends. After incubation, the cells were fixed with 4 % PFA for 60 min followed by quenching of the aldehyde groups with glycine or ammonium chloride and permeabilization with saponin (0.1 %). Primary antibodies used were: rat anti-LAMP-1 and 2, goat anti-EEA-1, anti-cathepsin D and anti-Rab3, mouse anti-Rab7, rabbit anti-Arl8b, mouse anti-CD63, rabbit anti-VAMP7 was made in-house by A. Peden and polyclonal guinea-pig anti-ADRP antibody. Cells were then incubated with the primary antibodies for 1 h at RT, washed, and finally incubated for another 1 h or overnight with the secondary antibodies conjugated with fluorophores. Antibody dilution was 1:500 for secondary antibodies conjugated with Cy3 or 1:200 for secondary antibodies conjugated with Cy5, Cy2 and Alexa 488. DAPI was used to visualize nuclei. Neutral lipids were stained with Bodipy 493/503 for 1 h. FC and ChA were stained with filipin for 30 min at RT. Coverslips were mounted with mowiol/DABCO and imaged using a LSM710 confocal microscope with a 63x oil-immersion objective with 1.4 NA.

The images were randomly acquired and 7 different fields were imaged for each experimental condition. For a given staining the images were acquired with the same settings. In each field cells that were clustered were ignored in the analysis because it was difficult to identify their boundaries and define the contours of the organelles of interest. Confocal images were obtained from three independent experiments and in total more than 45 cells were analysed per condition. Quantification of the number and size of the lipid-rich structures and lysosomes were performed using ImageJ software. The backgound was subtracted from the total fluorescence intensity of each region of interest (ROI).

For lysosome positioning within the cells, cell membrane and nuclear contours were extracted manually using ImageJ selection tools. Lysosome location and coordinates were also detected by drawing the ROI of each lysosome. Cell and nuclear contours and lysosomes XY coordinates were exported for posterior analysis on a custom Matlab Script. The measurement axis for each lysosome was defined as a straight line that crossed the nucleus and the lysosome centroids. This line’s intersection with the nuclear membrane and the plasma membrane originated the final coordinates for the calculation. Distances were measured using the Euclidian distance between the nuclear membrane, the lysosome and the plasma membrane. All calculations and the appropriate ratios were calculated using Matlab. The results were binned and plotted by fraction of lysosomes per distance to plasma membrane.

To quantify colocalization between lysosomes (LAMP-2-positive) and the indicated proteins (GTPases and CD63) we measured the area overlap between their segmented objects. Before segmentation, the images were enhanced by applying a median filter. The LAMP-2 objects were manually outlined and binarized. The GTPases and CD63 objects were automatically identified using ImageJ Analyse Particles function after applying a threshold and binarization. The threshold method applied was the same for each protein in different experimental conditions. The boundary coordinates of both the objects and lysosomes were extracted, by using the Roi.getCoordinates function of ImageJ, to estimate the number and size of each object per lysosome. The position of GTPases and CD63 vesicular objects were intersected with that of LAMP-2 objects to obtain the ones that were within or intersecting the LAMP-2 boundaries.

For live imaging, cells were seeded in Lab-Teck from Nunc plate and lysosomes were labeled overnight with 50 µg/mL of dextran conjugated with Alexa 647. To ensure that dextran was incorporated within the lysosomes, the overnight pulse was followed by a 4 h chase.

All the information regarding the antibody and dyes are available in Supplementary Table I - III.

### Electron microscopy

For conventional EM, cells were seeded on glass coverslips and, after exposure to the desired lipids, were fixed in 2 % PFA/2 % glutaraldehyde in 0.1M phosphate buffer. Cells were then osmicated, treated with tannic acid, dehydrated, infiltrated with Epon and mounted on Epon stubs all as described ^45^. After polymerisation overnight at 60 °C coverslips were removed from Epon stubs with liquid nitrogen. 70 nm sections were cut en face and stained with lead citrate before examination on a Jeol 1010 transmission electron microscope and images acquired with a Gatan OriusSC100B charged couple device camera.

For immuno-gold EM, cells grown in flasks were fixed in 4 % of PFA in 0.1 M phosphate buffer and then infused with 2.3 M sucrose, supported with 12 % gelatin and 80 nm sections cut at −120 °C. Sections picked up in a 1:1 mix of methyl cellulose/2.3 M sucrose were labelled at room temperature (RT) with primary antibodies, bridging antibodies and protein A gold as described ^46^. Primary antibodies were cathepsin D and LAMP-1. Protein A gold (10 nm) was from the University of Utrecht.

### DQ-BSA degradation

DQ-BSA degradation was assessed as described ^37^. The cathepsin L activity was evaluated by normalizing Magic Red fluorescence to Alexa Fluor 647-dextran fluorescence labeled lysosomes. The cells were incubated with Magic Red cathepsin L substrate for 15 min, washed and imaged under a confocal microscope. As positive control, lysosomes were alkalinized with 100 nM of Bafilomycin (Baf A1) added 1 h before the cathepsin L-substrate.

### Lysosome pH measurement

In lipid treated cells, lysosome pH was measured after an overnight incubation with dextran conjugated with FITC (250 µg/mL) and dextran conjugated with Alexa Fluor 647 (50 μg/mL) as described elsewhere ^37^. To measure the fluorescence intensity ratio between FITC- and Alexa Fluor 647-dextran, lysosomes with positive signal for both green and red channel were individually isolated. After background removal the fluorescence intensity of the two channels was quantified by the ImageJ Fiji software (Coloc2 puglin).

### Evaluation of LAMP-2 at the plasma membrane by flow cytometry

Cells treated for 72 h with lipids were washed with HBSS and then incubated at 37°C with 4 mM CaCl2, in the presence or absence of 5 µM ionomycin, during 5 min. After incubation, cells were shifted to 4°C, washed with flow cytometer buffer (PBS supplemented with 1% FBS and 2 mM EDTA), incubated with rat anti-mouse LAMP-2 antibody (1:50 in flow cytometer buffer), for 30 min followed for another 30 min with a secondary Cy5-anti rat antibody (1:250). Before analysis, cells were also stained with 50 µg/mL propidium iodide (PI). Data were acquired in a BD FACScantoTMII flow cytometer (Becton Dickinson): PI fluorescence and LAMP-2 were measured in the PI channel (585/42 BP) and Cy5 channel (665/20 BP), respectively. The software for acquisition was BD FACSDIVA Diva, and data from at least 5000 cells were analysed with FlowJo. All PI-positive cells were excluded for the quantification.

### β-hexosaminidase release assays

The β-hexosaminidase (β-hex) release assay was performed as described ^47^. For this assay, RAW cells were treated for 72 h with lipids.

### Plasma membrane staining and FM4-64 internalization

Cells treated with lipids were incubated with 30 μg/mL FM4-64 in DMEM with HEPES, for 15 min on ice and then washed with PBS. Cells were then incubated with complete DMEM without phenol red and incubated at 37 °C for different time points. Live cells were imaged at 37°C under an LSM880 confocal system using airyscan fast with 63x/1.4NA objective. Z-stack images were acquired at time 0, 15 and 45 min after chase. Images were processed only by using airyscan processing with the default strength value. Quantification of the internalized dye was measured using ImageJ software, by calculating the ratio between the cytosolic fluorescence to the plasma membrane intensity. Values were normalized using the timepoint zero value as a reference.

### Immunoblotting

Cell lysates were prepared with RIPA buffer in presence of protease inhibitors. After clearance the protein concentration was determined using the PierceTM BCA Protein Assay Kit. Samples were then mixed with Laemli buffer, heated for 5 min at 95 °C and loaded (40-70 µg) on 10-15 % SDS polyacrylamide gels. After electrophoresis, proteins were transferred at 4 °C onto activated PVDF membranes in transfer buffer for 2 h at 300 mA with gentle agitation. Membranes were then blocked with blocking buffer for 1 h at RT followed by an overnight incubation at 4 °C with the following primary antibodies diluted in blocking buffer: mouse anti-LAMP-2 and anti-LAMP-1, Arl8b, CD63, VAMP7 (was made in-house by Andrew A. Peden) and anti-Rab3, anti-calnexin, cathepsin D and Rab7a. After incubation with primary antibodies, membranes were washed with TBS Tween and incubated for 1 h at RT with the corresponding horseradish peroxidase secondary antibody diluted in blocking buffer. Blots were visualized using ECL Prime Western Blot Detection reagent (GE Healthcare) and a Chemidoc Touch Imaging System (Bio-Rad Laboratories). Bands were quantified using Image Lab 6.0 software from Bio-Rad.

### Quantitative RT-PCR

Total RNA was extracted with NZY total RNA isolation kit and reverse transcription was performed using NZY first-strand cDNA synthesis kit. Quantitative PCR was performed in a 96-well plate using the SYBR green master mix using AB7300 Real-Time PCR thermal cycler with Step One software (v2.2.2; Applied Biosystems). *Hprt* and *Pgk1* were used as housekeeping genes to normalize the expression. Target gene expression was determined by relative quantification (ΔΔCt method) to the housekeeping reference gene and the control sample. Hprt primers were from QIAGEN, the other primer sequences are indicated in Supplementary Table IV.

### Statistical analysis

Difference between mean lipid concentration and the ± standard error of the mean (SEM) were calculated in R. The ‘ggplot2’ package was used for boxplot. Robust statistical tests like *Kruskal*-*Wallis* test followed by unpaired two-samples Wilcoxon test were used to assess statistical differences in lipid concentration between patients groups. Remaining results are expressed as the mean ± SEM. Statistical significance was assessed using GraphPad software by one way ANOVA followed by *Kruskal*-*Wallis* test with Dunn’s multiple comparisons, two ways Tukey post-test or Student t-test depending on data distribution. A *p* value of 0.05 was considered to be statistically significant.

## RESULTS

### Cholesteryl hemiazelate is the most prevalent cholesteryl hemiester in plasma of CVD patients

One of the major modifications of LDL in the intima is oxidation of their polyunsaturated fatty acid esters of cholesterol which results in the accumulation of so-called “core aldehydes” ^48-50^ in the resulting fatty streaks and atheromata. As we have argued eslsewhere ^35^, the oxygen-rich environment in the arteries promotes the conversion of “core aldehydes” to ChE that could be one of the important etiological agents in atherogenesis. We previously investigated this possibility by studying the conversion of a macrophage cell line (RAW 264.7 cells, hereafter referred to as RAW cells) to “foam cells” upon exposure to ChS, a commercially available but biologically irrelevant ChE ^36, 37^. ChE, being considerably more polar amphiphiles than FC, are expected to partition from the oxidized lipid core of atheromata into the plasma. We, therefore, determined the plasma levels of ChE in CVD patients, by applying quantitative mass spectrometry-based shotgun lipidomic analysis.

In order to obtain reliable quantification of ChE, we infused the synthetic standards ChS, ChG (the cholesteryl hemiester of glutaric acid), and ChA and observed that this class of lipids ionize efficiently in the negative ion mode as deprotonated anions. Moreover, we were also able to establish the MSMS fragmentation of the different ChE species, their unambiguous detection and quantification (Supplementary Figure II A-C). We then assessed whether ChE could be detected and quantified in plasma samples. Increasing amounts of a test plasma sample were extracted together with 100 pmol of ChS, an internal standard which is not normally a component of plasma, and looked in the high resolution (HR)MS and MSMS spectra for the ChE mass spectrometric “fingerprint”. Different ChE, with differing carbon lengths and degrees of unsaturation were observed. Most importantly, their intensities were correlated with the sample volume (Supplementary Figure III). The dynamic range and limits of quantification were established by “spiking” increasing amounts of ChS in 2 µL of the plasma sample (Supplementary Figure IV). The dynamic range was 500 and the inferior limit of quantification was 0.5 µM.

In our plasma samples, two ChE were quantitatively analysable (concentrations >0.5 µM): ChA and a cholesteryl hemiester of 1,11-undecendioic acid. ChA was quantifiable in 88.8 %, 67.0 %, and 36.5 % of acute coronary syndrome (ACS), stable angina pectoris (SAP), and control cohorts, respectively (Figure 1A). The cholesteryl hemiester of 1,11-undecendioic acid was only quantifiable in 67.0 %, 31.7 %, and 11.5 % of the ACS, SAP, and control cohorts, respectively (Figure 1B). Thus, ChA was the most prevalent of the two ChE. ChA concentrations were significantly higher in ACS and SAP when compared to age-matched controls. The mean differences from control were 1.04 ± 0.15 µM for ACS and 0.73 ± 0.16 µM for SAP. Futhermore, the highest ChA concentration was 5.95 µM (Figure 1A). This finding is well correlated with previous work from other laboratories showing that cholesteryl-9-oxononanoate (the precursor of ChA) was the principal “core aldehyde” in oxidized LDL ^48, 51^ and in atheromata ^49^. We believe that the second quantifiable ChE observed (cholesteryl undecenedioic acid hemiester) is also formed from cholesteryl linoleate but this will require appropriate chemical verification. Since the ChA relevance in the context of CVD has never been investigated, we next investigated the atherogenic properties of this lipid.

**Figure 1:**
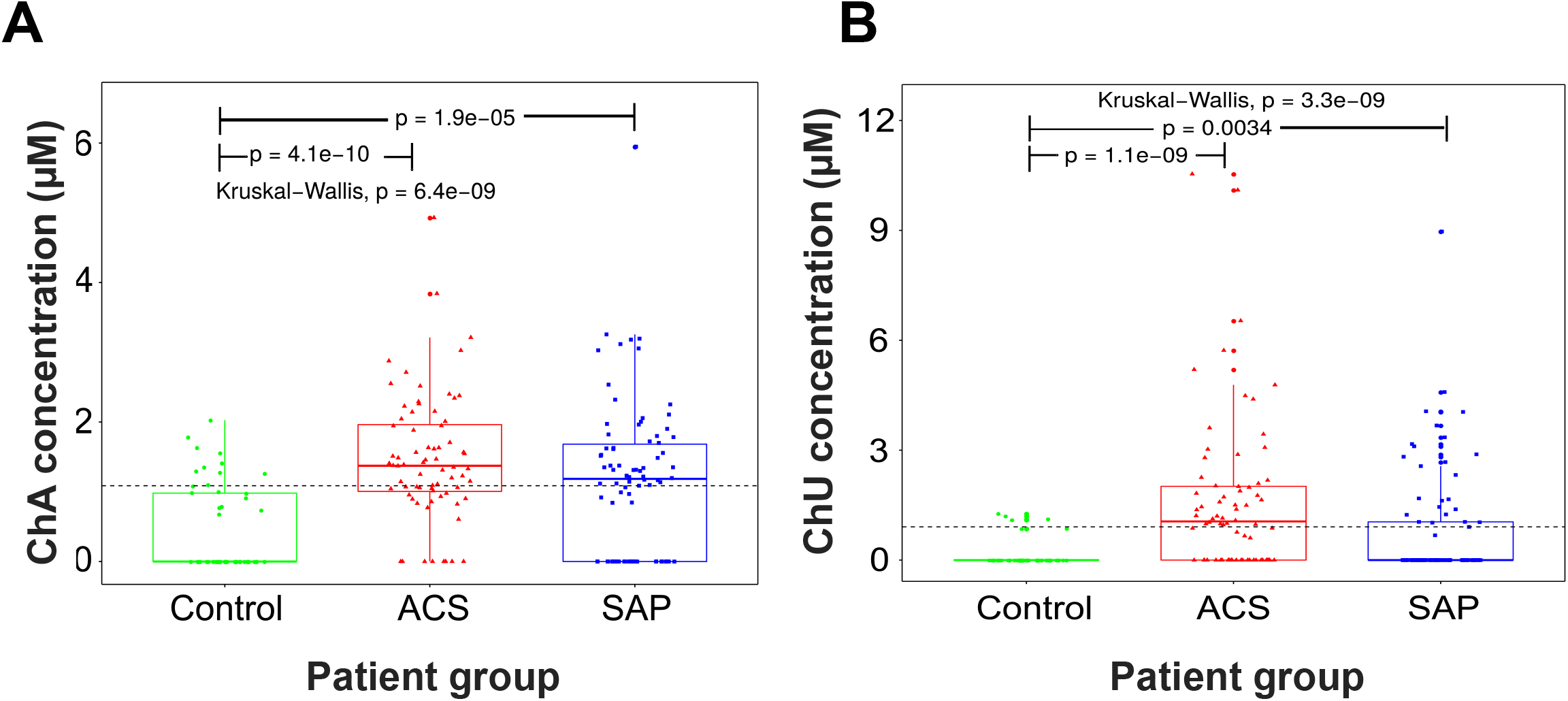
ChA levels are increased in plasma of CVD patients. Boxplots depicting ChA (**A**) and Cholesteryl hemiester of 1,11-undecendioic acid (ChU) (**B**) concentrations across patient groups. Kruskal–Wallis one way analysis of variance was applied to test whether ChA concentration across samples originate from the same distribution. Unpaired two-samples Wilcoxon test was applied as a post hoc test to evaluate significant differences in lipid levels for a patient group versus the control group (indicated by “p”). The horizontal black dashed line indicates the global mean across all samples. Plasmatic concentration of ChA was obtained by shotgun lipidomics of 72 donors with acute coronary syndrome (ACS, including ST-elevation and non-ST-elevation myocardial infarction and unstable angina pectoris), 82 donors with stable angina pectoris (SAP) and 52 age-matched control cohort.

### Cholesteryl hemiazelate is toxic to macrophages and induces lipid accumulation and formation of peripheral enlarged lysosomes

Macrophages are highly relevant in the initiation and development of atherosclerosis. Since atherogenesis is an extremely slow process and it is not known how long macrophages have to be exposed to the toxic products in modified LDL to develop the “foam cell” characteristics and, eventually, die; it was decided that this first study on the effect of ChA towards macrophages should preferably be done using an “eternalized” macrophage cell line (RAW cells) rather than primary cells. Due to the poor solubility of ChA in the cell culture medium and to avoid artefacts due to possible uptake of microcrystals of ChA, all experiments were performed with ChA:POPC liposomes (65:35, molar ratio). POPC liposomes were used as control. Earlier work from our laboratory ^36^ had shown that POPC liposomes are almost as efficient as vehicles for ChE as native LDL or acetylated-LDL.

The toxicity curve for ChA on RAW cells is shown in Figure 2A. Over a 72 h exposure time, the LD50 is 2000 µM and is preceded by a wide (starting at about 3 µM ChA) hormetic zone. In order to obtain data representing both pre-toxic and toxic phenomena within practically convenient exposure times we used a ChA concentration slightly below the measured LD50 value, namely 1500 µM. At a first glance, and considering that ChA concentrations in plasma of CVD patients is only about 1.5 µM on the average (Figure 1A, dashed line), the ChA concentrations used here might seem very high. However, taking into account that foam macrophages are formed in fatty streaks or atheromata in the arterial intima, the concentration of ChA that a macrophage is exposed during atherogenesis is much higher than that found in circulation. Since ChA is an amphiphilic molecule closely related to cholesterol, is expected a strongly partition to the surface of atheromata, thus increasing the local concentration. From earlier work reported from one of our laboratories^52-55^ the lipid phase/aqueous phase partition coefficient for amphiphiles similar to ChA is expected to be about 10^5^ for more polar molecules like lyso-PC and greater than 10^6^ for more apolar amphiphiles like dehydroergosterol, a cholesterol analog. The concentration of ChA detected in plasma represents its value after the equilibration with all cell surface membranes directly in contact with the blood. The ChA mean value of 1.5 μM found in the plasma of ACS and SAP patients is expected to be the result of the partition between the aqueous phase, the serum albumin, and the polar lipid monolayers at the surface of colloidal lipid particles (the lipoproteins). Considering that the rate constants for the ChA insertion and desorption to the lipoprein surface monolayer, the association and dissociation with serum albumin, and the equilibrium constants for association with both the lipoprotein surfaces and albumin are known, the exact concentrations of ChA associated with the lipopreins, albumin and free in the plasma aqueous phase can be calculated. These values are not known specifically for ChA but the values for a large number of amphiphiles have been previously measured by our group. Taking this into accont, the concentration of ChA detected in the plasma of ACS patients in the polar lipid monolayer that constitutes the surface of the plasma lipoprotein particles yields a value of 0.56 M (see Supplementary Table V). Thus, given a concentration of ChA in plasma of 1 µM we may expect its concentration in the atheromatal lipid mass to be on the order of ∼0.1 M or higher ^55^. This is the effective “physiological concentration” that a macrophage in atheroma is exposed to.

**Figure 2:**
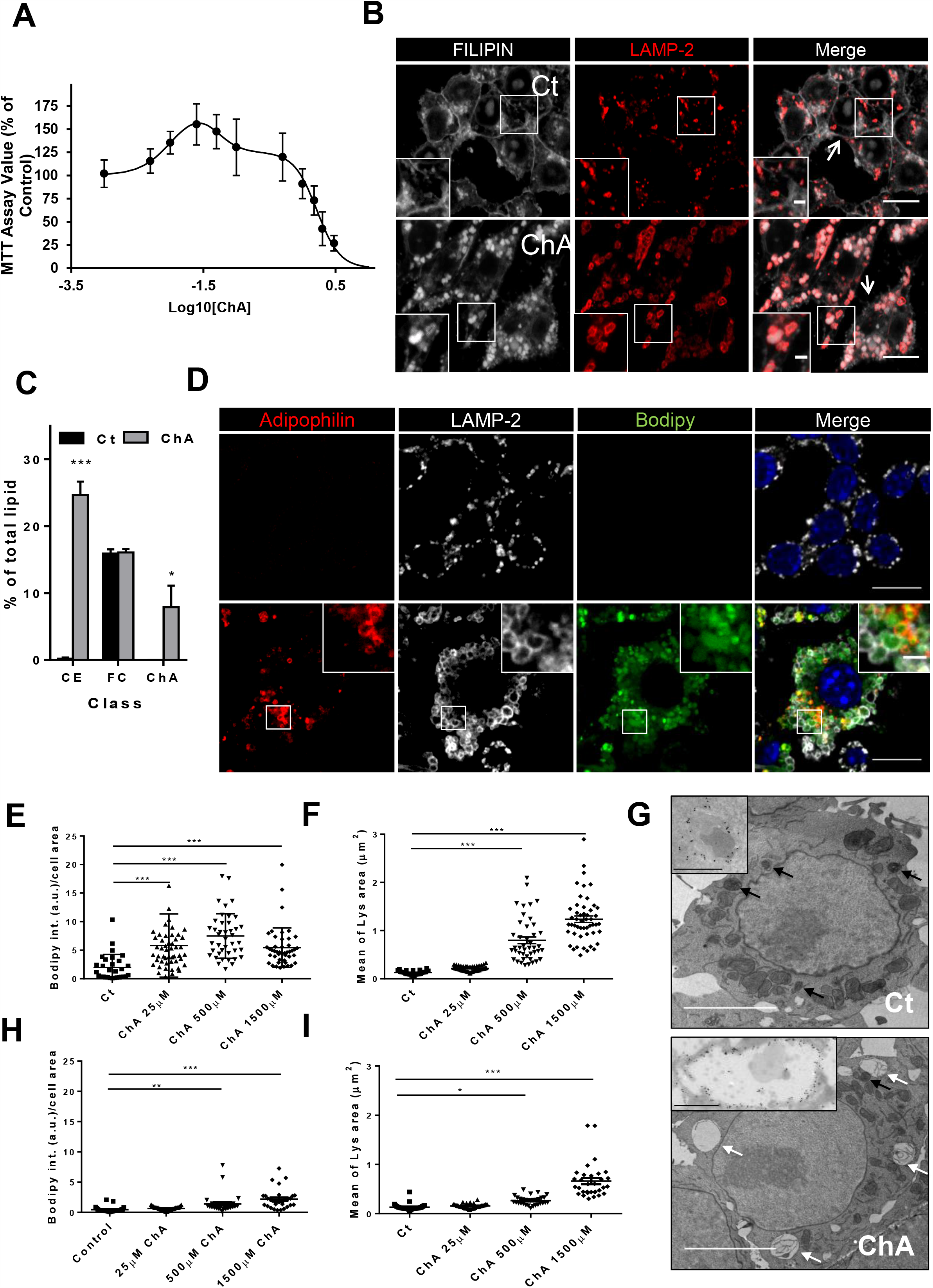
ChA is toxic and induces lysosome enlargement and lipid accumulation in macrophages. **A**. Effect of ChA on the viability of RAW cells. RAW macrophages were incubated with ChA or with POPC (control cells) during 72 h and cell viability was evaluated by the MTT assay. The results are present as mean ± SEM of four independent experiments. The line is a theoretical fit using the Hill Equation. **B**. Representative images of RAW macrophages incubated with 1500 µM of ChA or with POPC (control cells, Ct) for 72 h. Filipin and Lysosomal Associated Membrane Protein-2 (LAMP-2, a marker of late endocytic compartments) staining of control and ChA-treated cells. The merged images show ChA in white and lysosomes in red. The insets are enlargements of the areas outlined with the white boxes. Arrows point to plasma membrane of RAW cells. **C**. Quantification of cholesteryl esters (CE), free cholesterol (FC) and ChA by shotgun lipidomics MS analysis of the total lipid extracts of cells incubated with ChA or POPC for 72 h. The results are mean ± SEM of three independent experiments. The p values were obtained by two way ANOVA Sidak test ***, p<0.001; *, p<0.05. **D**. Representative images of adipophilin, LAMP-2 and BODIPY. Nuclei were labeled with DAPI. The merged images show lipid droplets in red, lysosomes in white, neutral lipids in green and nuclei in blue. The insets are enlargements of the region outlined in the main images. Scale Bars, 10 μm in the main images and 2 μm in the insets. **E and F**. Quantification of neutral lipid accumulation (**E**) and lysosome (Lys) area (**F**) in macrophages pulsed for 72 h with POPC or ChA. The results are mean ± SEM of at least 30 cells in total (n=3). **G**. Transmission electron microscopy of RAW cells treated for 72 h with POPC or ChA liposomes. ChA-treated cells exhibit larger lysosomes as confirmed by LAMP-1 immuno-gold staining (insets). Electron dense lysosomes and enlarged electron lucent lysosomes are indicated by black and white arrows respectively. Scale bars, 500 nm and the insets 200 nm. Quantification of neutral lipid accumulation (**H**) and lysosome area (**I**) in RAW cells pulsed for 72 h with POPC or ChA followed by a chase of 72 h. The results are mean ± SEM of at least 30 cells in total (n=3). In **E, F, H** and I, the p values were obtained by one way ANOVA-Kruskal-Wallis test ***, p<0.001; **, p<0.01; *, p<0.05.

Considering our previous work ^36, 37^ with ChS, we enquired whether ChA was also targeting the lysosomes in RAW cells. Lipid accumulation in late endosomes and lysosomes (LE/Lysosomes, hereafter collectively referred to as lysosomes, unless stated otherwise) as well as lysosome morphology were assessed by confocal microscopy and electron microscopy. RAW cells were incubated with increasing amounts of ChA for 72 h and then fixed. Lysosomes were visualized with antibodies against the lysosome associated membrane protein-1 or 2 (LAMP), that in the steady state is mostly confined to the limiting membrane of lysosomes, by immunofluorescence (IF). ChA and neutral lipid accumulation was observed by staining the cells with Filipin and Bodipy, respectively. Filipin besides being a stain for FC also stains ChS ^37^ and ChA (Supplementary Figure V A). Figure 2B showed that in ChA-treated cells Filipin staining was more intense in the enlarged LAMP-2 vesicles, a protein that is mostly confined to the limiting membrane of lysosomes, than in the plasma membrane (PM). In control cells, Filipin, stained mainly the PM. To conclude that the lysosomal Filipin staining was due to ChA accumulation rather than to FC, the levels of these lipids were quantified, by shotgun lipidomics, in lysates of cells incubated with ChA and POPC (Figure 2C). The data showed that while FC levels remained identical in ChA- and POPC-treated cells, the Choleseryl ester (CE, neutral lipids) and ChA levels were only increased in cells exposed to ChA. This result led us to assume that Filipin staining in lysosomes was mainly due to ChA accumulation rather than to FC (Figure 2B). The higher concentrations of CE in ChA-treated cells also suggested that some ChA hydrolysis occurred followed by re-esterification and intracellular storage in cytosolic lipid droplets. To confirm this, the intracellular neutral lipid distribution (as seen by Bodipy staining) was compared with that of LAMP-2 and adipophilin (lipid droplet marker). Quantification of total Bodipy intensity (Figure 2E-F) revealed an increase in neutral lipids in ChA-treated macrophages.

Furthermore, the brightest spots for neutral lipids staining were surrounded by adipophilin indicating that they were cytosolic lipid droplets (Figure 2D, lower panels and enlarged regions outlined by the squares) confirming partially ChA hydrolysis. Confocal images also showed that macrophages incubated with ChA exhibited some neutral lipid accumulation inside enlarged lysosomes. This outcome is probably a consequence of lysosome malfunction culminating in accumulation of undegraded endocytosed neutral lipds that exist in the cell culture medium.

Next, the lysosome area as a function of ChA concentration was quantified (Figure 2F). After an incubation of 72 h with ChA, RAW cells exhibited a significant increase in lysosomal area (0.8 ± 0.07 µm^2^ and 1.2 ± 0.068 µm^2^ for 500 and 1500 µM ChA, respectively) in comparison to control cells (0.13 ± 0.01 µm^2^) in a dose-dependent manner (Figure 2F, n = 50). To gain more insights at the ultrastructural level, we also performed TEM and immuno-gold staining to visualize lysosomes and LAMP-1 distribution, which in control and ChA-treated cells presents the same distribution as LAMP-2 (Supplementary Figure V B). Whilst in control cells lysosomes had a typical electron dense appearance, many of the lysosomes in ChA-treated macrophages were enlarged and more electron lucent (Figure 2G).

To ensure that the phenotypes described above for ChA were specific for this lipid, RAW cells were also exposed to FC:POPC liposomes at a molar ratio 65:35 and to cholesteryl linoleate (ChL):POPC emulsions as controls (Supplementary Figure V C). After 72 h incubation with both lipids, RAW cells also exhibited cytosolic neutral lipid accumulation but less exuberantly than with ChA:POPC treated cells. Interestingly, no changes in lysosome morphology or lipid accumulation within these organelles were observed, indicating that the phenotypes observed in ChA-incubated cells were specific for this lipid.

Previous work with ChS in our laboratory showed that endolysosomal enlargement and lipid accumulation were irreversible^36^. We, therefore, questioned if ChA was also inducing a permanent effect. Lysosome area and lipid accumulation irreversibility was assessed through a pulse-chase experiment. Briefly, after 72 h of ChA treatment, macrophages were incubated in a ChA-free medium for an additional period of 72 h (Figure 2H and 2I). The results indicate that the intracellular lipid accumulation (Figures 2E and 2H) and lysosome area (Figures 2F and 2I) induced by ChA were, to some extent, partially reversible. These results suggest that the intracellular hydrolysis of ChA is more efficient than the hydrolysis of ChS.

### In ChA-treated macrophages lysosomes are dysfunctional

It is expected that changes in lysosome morphology will affect intracellular trafficking and their degradative capacity^56^. Thus, in order to evaluate if ChA was affecting the endocytic pathway, we started by challenging cells with different endocytic cargos and then their intracellular transport was followed. The results suggested that the uptake of endocytic cargo by macrophages, whether receptor-mediated or not, was not affected by ChA (Supplementary Figure VI A-F). However, we observed that transferrin recycling and cargo transport up to lysosomes were delayed (Supplementary Figure VI E-I and Supplementary Material). Since lysosomes play a central role in both receiving and degrading macromolecules from the endocytic membrane-trafficking pathways, we next examined the effect of ChA in cargo degradation. For this purpose, we used DQ-BSA, a fluorogenic substrate for lysosomal proteases. Enzymatic cleavage of DQ-BSA produces highly fluorescent products. The results shown in Figure 3A showed a higher enzymatic cleavage of DQ-BSA in control cells than in cells exposed to ChA. Furthermore, the rate of degradation of DQ-BSA by the lysosomal proteases was significantly slower in cells exposed to ChA in comparison to control cells, suggesting that the catalytic activity of this organelle was affected (Figure 3B). However, since we observed a delay in cargo delivery to lysosomes in ChA-treated cells, the decrease in DQ-BSA degradation can be affected by this defect in the endocytic pathway. To circumvent this problem, we measured the activity of lysosomal cathepsin L through a fluorogenic assay using the “Magic Red” cathepsin substrate (Figure 3C and 3D). Bafilomycin A (Baf A1), a vacuolar-type proton-ATPase inhibitor which increases lysosomal pH was used as control. For this set of experiments lysosomes were previously labeled with Alexa 647-dextran, which fluorescence is not pH-dependent, followed by addition of the Magic Red substrate. Cathepsin-L proteolytic activity was quantified by ratiometric fluorescence (Figure 3C; pseudocolor scale). The enlarged lysosomes in ChA-treated cells presented a lower ratiometric value than the control cells (Figure 3D), indicating a reduction in proteolytic activity. Thus, besides the delay in lysosomal cargo delivery, enlarged lysosomes also present a defect in cargo degradation suggesting that the LAMP-2-vesicles are indeed non-functional lysosomes.

**Figure 3:**
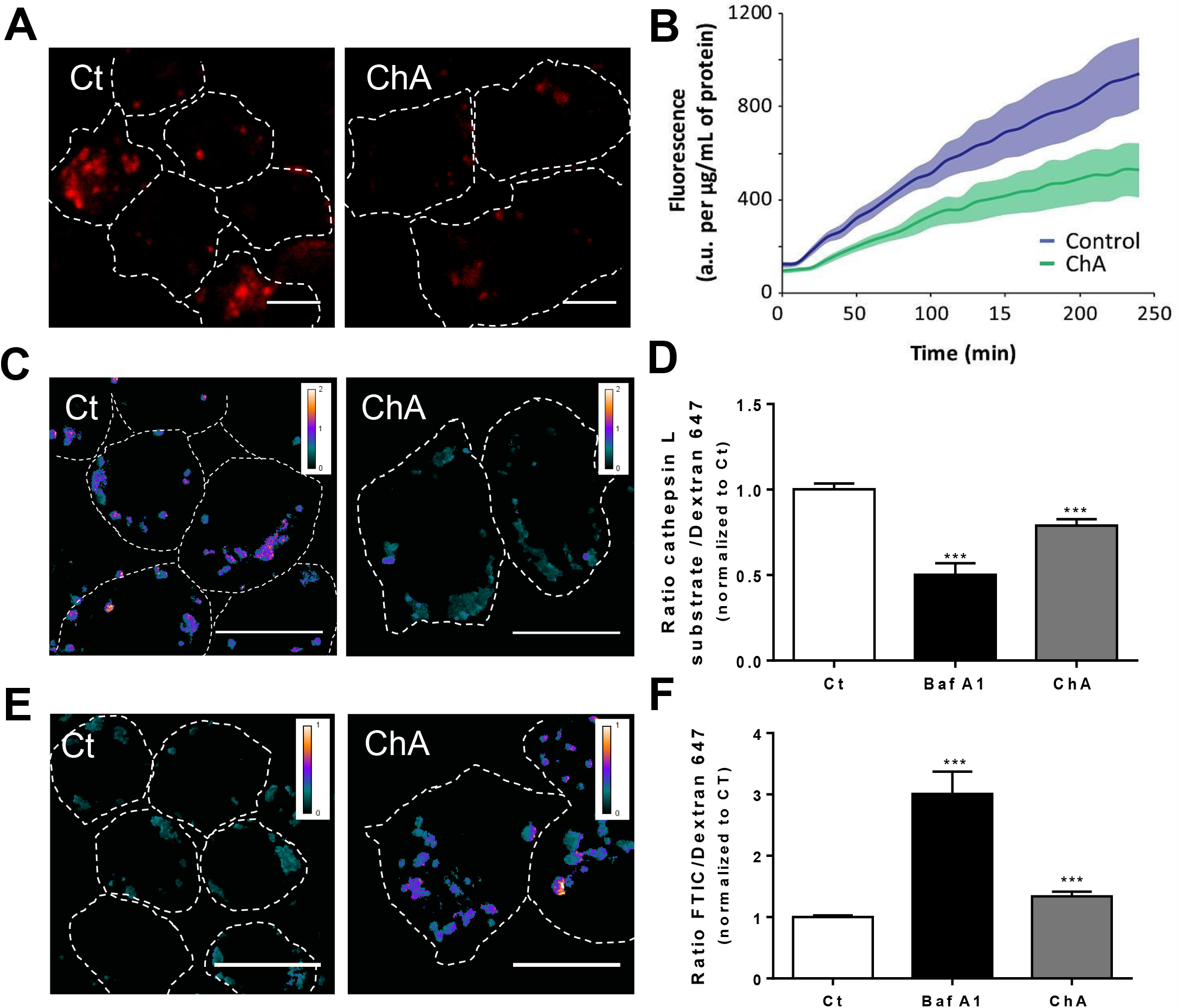
ChA induces lysosome dysfunction. RAW cells were incubated with 1500 µM ChA or with POPC (control cells, Ct) for 72 h. **A**. Representative images of DQ-BSA degradation in control and ChA-treated cells. Cells were loaded for 3 h with DQ-BSA, chased 4 h to allow cargo degradation and then imaged using confocal microscopy. **B**. Time-course of DQ-BSA degradation in RAW cells. Fluorescence was measured using a microplate fluorescence reader. The data represent the mean ± SEM (n = 3). SEM is represented by the color area around the mean line. **C**. Representative live cell images of lysosomal degradative capacity of cathepsin L substract. After cells treatment, lysosomes were labeled with Alexa Fluor 647– dextran and then incubated for 15 min with Magic Red cathepsin L substrate. Cells were imaged by confocal microscopy. The representative images show the ratio between Magic Red and Alexa Fluor 647–dextran fluorescence. **D**. Cathepsin L activity was quantified in control and ChA-treated cells by normalizing Magic Red fluorescence to Alexa Fluor 647– dextran fluorescence. As positive control, lysosomes were alkalinized with bafilomycin (Baf A1) added before Magic Red. **E**. Representative images of RAW cells obtained by confocal microscopy after incubation with FITC-dextran, a pH-sensitive dye and Alexa Fluor 647-dextran, a pH-insensitive dye. The images show the fluorescence ratio between the two dextrans. **F**. Quantification of lysosomal pH in live cells measured by ratiometric fluorescence. As described in **D**, Baf A1 was used as positive control. In **A, C** and **D**, dashed lines show the edges of Raw cells. Three independent experiments were performed and each time 20 cells were analysed. Error bars represent SEM. ***, p<0.001 determined by one way ANOVA-Kruskal-Wallis test. Scale bars, 5 μm.

The proteolytic activity of lysosomal enzymes is strictly dependent on pH values in the lysosomal lumen. Therefore, a slight variation in lysosomal pH can compromise the degradative capacity of these organelles. Thus, we decided to assess whether lysosome luminal pH, in ChA-treated cells, was higher when compared with control cells. Lysosomal pH was measured in cells previously incubated with FITC-(Fluorescein isothiocyanate, sensitive to pH) and Alexa 647-dextran by quantifying the ratio between the emission at 488 and 645 nm. RAW cells treated with Baf A1 were used as positive control. The results presented in Figure 3E and 3F show that ChA increased the lysosome pH, which can explain the loss of degradative capacity of these organelles.

### ChA drives lysosome biogenesis in macrophages

Lysosomes are able to sense their content in stress conditions and regulate their own biogenesis by transcriptional encoded mechanisms. It has been demonstrated that induction of lysosomal stress with potent lysosomal inhibitors, such as chloroquine, or atherogenic lipids, such as cholesterol crystals, in macrophages leads to the upregulation of genes involved in autophago-lysosome formation and acid hydrolases, important in the degradation of macromolecules ^26, 57^. The master regulators of lysosome biogenesis are the transcription factors EB (TFEB), the related Transcription Factor Binding To IGHM Enhancer 3 (TFE3) and Melanocyte Inducing Transcription Factor (MITF), which engage a gene network containing >400 genes, many of which encode lysosomal proteins ^57, 58^.

To assess if lysosome biogenesis was occurring, in ChA-treated macrophages, we started by investigated TFEB, TFE3 and MITF mRNA expression by qRT-PCR experiments (Figure 4A). The time-course data suggested that these transcription factors respond to ChA and the highest levels of transcription occur between 12 and 24 h. These transcription factors control the expression of the coordinated lysosomal expression and regulation (CLEAR) genes. For the lysosomal genes evaluated, we observed an increase in their transcription after 12 h. *LAMP-1* expression started to increase at 12h and was sustained with time and, *lipa* peaked at 24 h (Figure 4B). To further investigate these results, we examined if the enhanced mRNA levels corresponded to an increase in lysosomal protein levels and in lysosome number. As shown in the Figure 4C-F there was an increase for LAMP-1, LAMP-2 and Pro-Cathepsin D at 72 h incubation time. For LAMP proteins the increase was significant and dose-dependent. Strinkingly, this data is well correlated with an elevation in the number of lysosomes per cell, indicating that ChA induces *de novo* lysosome biogenesis (Figure 4G). The increase in lysosome biogenesis and the transcriptomic activation of the lysosome pathway could compensate lysosome dysfunction as well as the absence of effects at the autophagy level. Indeed, besides the increase in *lc3* and *p62* mRNA levels observed in ChA-treated cells (Figure 4B) the autophagic flux and p62 protein levels showed no statistically differences when compared with control cells (Supplementary Figure VII).

**Figure 4:**
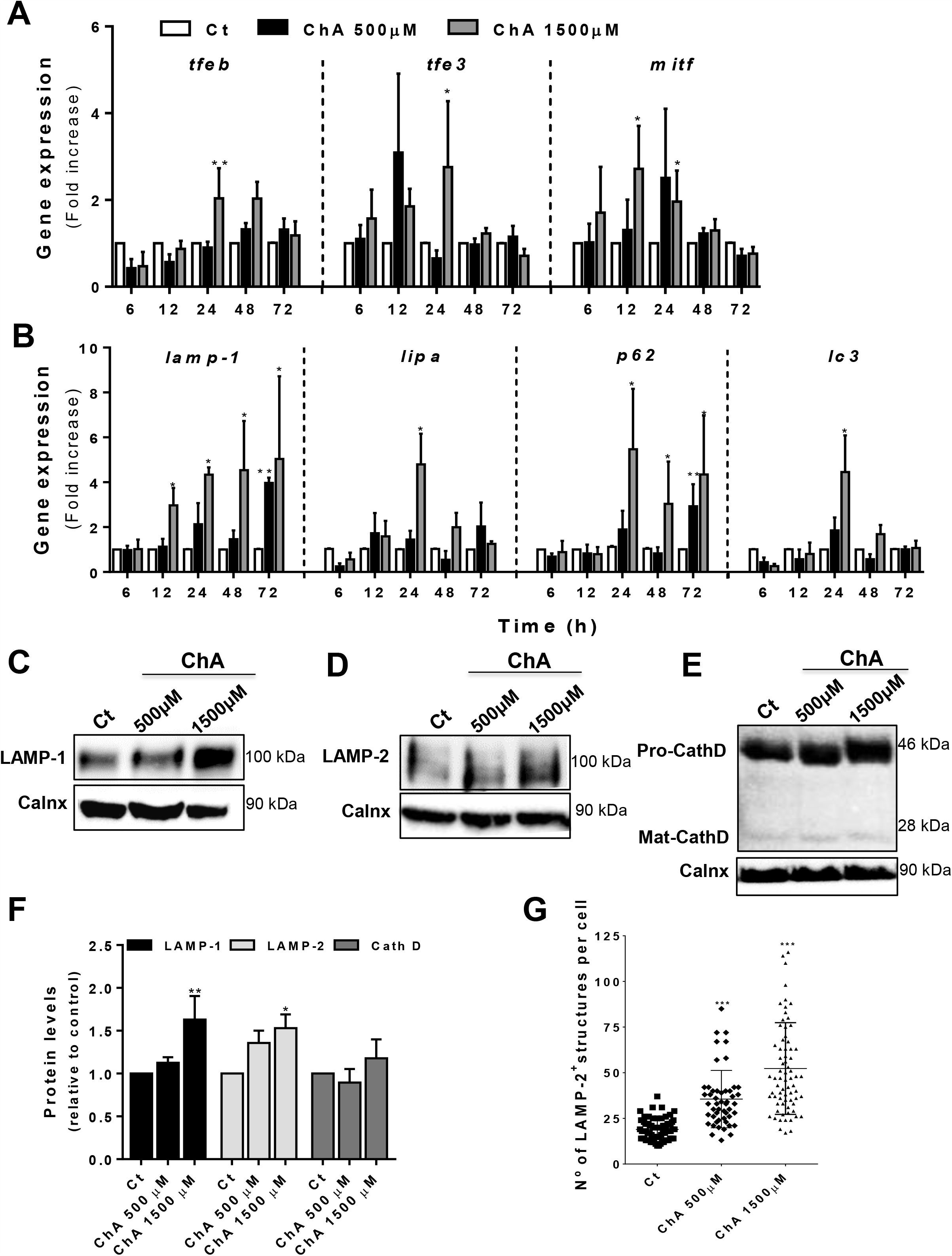
Lysosome dysfunction induces *de novo* lysosome biogenesis. **A**. Gene transcription levels of the three main transcription factors involved in lysosomal biogenesis – TFEB, TFE3 and MITF in function of the incubation time and lipid concentration. **B**. mRNA expression levels of *LAMP-1, lipa, p62* and *lc3* genes. mRNA levels were assessed by qRT-PCR. Data were normalized to the endogenous *hprt* and *pgk1* genes. The values are mean ± SEM of three independent experiments. The p values were obtained by one way ANOVA-Kruskal-Wallis test **, p<0.01; *, p<0.05. **C-E**. Western-blots showing LAMP-1 (**C**), LAMP-2 (**D**) and Cathepsin D (**E**) protein levels in lysates of cells treated with POPC (Ct) or ChA. **F**. Ratio between LAMP-1, LAMP-2, total Cathepsin D and Calnexin (used as loading control) of the quantified bands in western-blots. **G**. Quantification of total number of lysosomes per macrophages after lipid treatment for 72 h. In total, at least 50 cells were analysed per condition. ***, p<0.001**, p<0.01; *, p<0.05 determined by by one way ANOVA-Kruskal-Wallis test.

### In ChA-treated cells lysosomes are shifted towards the cell periphery

Lysosomes are considered a highly dynamic and heterogeneous organelle in their pH ^56^, subcellular position ^47, 59, 60^ and morphology ^61-63^. Increased TFEB levels were correlated with an increase in lysosome population near the PM ^64^. Taking this into account, we next investigated the effect of ChA on lysosome positioning, through an in-house produced bespoke analysis software, produced in house. As schematically illustrated in Figure 5A, the radial distance from the nuclear envelope to lysosomes was assessed in control and ChA-treated cells and normalized to the corresponding radial distance from the nuclear envelope to the PM. Thus, values close to zero are an indication of a more perinuclear localization and values near to 1 indicate a closer proximity of lysosomes to the PM. In Figure 5B, it is seen that 50 % of the total analysed lysosomes were at a normalized distance >0.7 in ChA-treated cells whereas in the control cells >50 % were at a normalized distance <0.5 from the nuclear membrane. Thus, lysosome distribution is considerably more peripheral in ChA-treated than in control cells.

**Figure 5:**
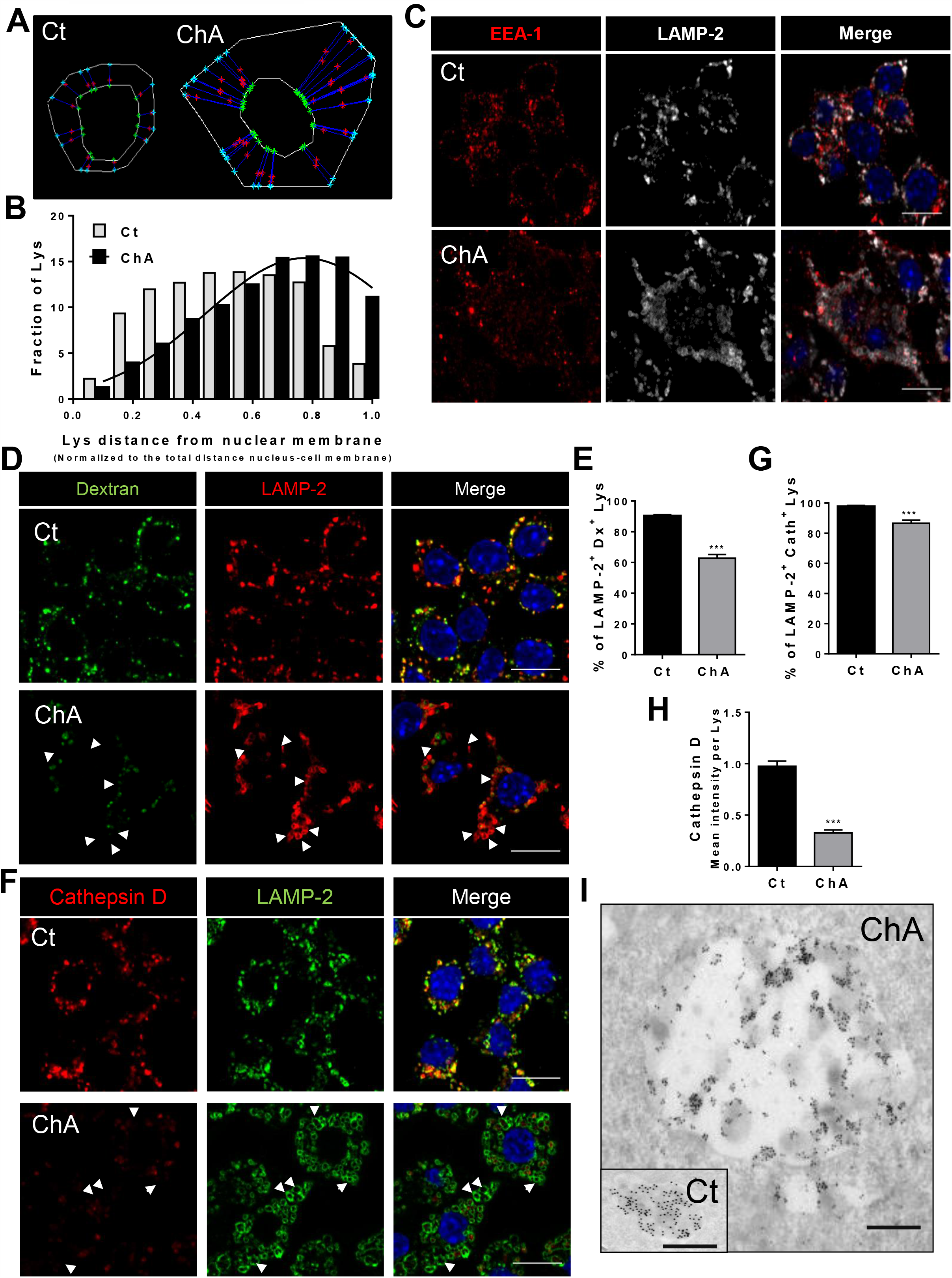
ChA induces changes in lysosome positioning and composition. RAW cells were incubated with 1500 µM ChA or with POPC (control cells, Ct) for 72 h. **A**. Schematic presentation of the methodology used to measure lysosome positioning in RAW cells treated with POPC or ChA, taking into account the difference in cell size. **B**. Shows the fraction of total LAMP-2-positive organelles as a function of distance from nuclear membrane. The distance between each lysosome and the nuclear membrane was performed using an in-house developed software and normalized to the total distance between nucleus and the plasma membrane. In ChA-treated cells the results were fitted with a Gaussian curve showing a markedly peripheral lysosome distribution. At least 45 cells were analysed from 3 independent experiments. **C**. Confocal images showing the intracellular distribution of Early Endosomes Antigen-1 (EEA-1, in red) and LAMP-2 (in white) in control (Ct) and ChA-treated cells. **D**. Confocal images showing the intracellular distribution of vesicles labeled with Alexa Fluor 647–dextran and LAMP-2. Arrowheads point to LAMP-2-positive vesicles that are not positive for dextran. **E**. Quantification of LAMP-2- and Alexa Fluor 647–dextran-positive vesicles. **F**. Representative images of RAW cells stained for LAMP-2 and cathepsin D. Arrowheads indicate LAMP-2-labeled organelles with no detectable cathepsin D. **G**. Quantification of LAMP-2- and cathepsin D-positive vesicles. **H**. Fluorescence intensity of cathepsin D per lysosome area. Data presented is the mean ± SEM of three independent experiences. In **E, G** and **H**, 15 cells per experiment were analysed. The results are mean ± SEM of 50 cells in total (n=3). Statistical significance was assessed by a t-test: ***, p<0.001. Scale bars, 10 μm. **I**. Immuno-gold EM showing Cathepsin D staining in Ct and in ChA-treated cells. Scale bars, 500nm.

Further characterization showed that the peripheral enlarged-lysosomal LAMP-2-positive vesicles were not positive for EEA-1 (Figure 5C). Thus, these large LAMP-positive organelles were not hybrid organelles of the endocytic compartment. Next, we labeled lysosomes by a pulse-chase experiment with fluorescent dextran. The results demonstrated that the percentage of fluorescent LAMP-2-positive endocytic compartments positive for dextran was 67.7 ± 1.7 % in ChA-treated cells and 90.4 ± 0.8 % in control cells (Figure 5D and 5E). However, the percentage of lysosomes identified as LAMP-2 and Cathepsin D double-positive organelles in ChA-treated cells was only slightly lower than that in control cells, 86.6 ± 2.0 % *vs* 97.7 ± 0.7 %, respectively (Figure 5F and 5G). Due to the larger area of these organelles in ChA-treated cells, the cathepsin D intensity per lysosome was lower than in lysosomes of control cells (Figure 5H). These results were in agreement with immune-gold-EM images for cathepsin D (Figure 5I), where the density of immune-gold particles per enlarged lysosome is reduced when compared with control lysosomes. Altogether, these data indicate that in ChA-treated RAW cells most of the enlarged lysosomes have cathepsin D though more than 30 % cannot be labelled by dextran suggesting that the sub-populations of LAMP-2-labeled organelles were different in ChA-treated and in control cells.

Some of the trafficking machinery involved in lysosome peripheral transport and positioning has been identified such as Arl8b and Rab7a GTPases. Arl8b GTPase is recruited by a multisubunit protein complex BORC to the lysosomal membrane recruiting kinesin-1 via SKIP, which catalyses lysosome movement to the periphery ^65^. Rab7a GTPase links lysosomes to dynein via RILP catalysing the lysosome movement to the perinuclear region ^66^. However, Rab7a can also link lysosomes to kinesin-1 via its effector FYCO1 translocating lysosomes to the periphery ^67^. Once at the periphery, Rab3a GTPase is necessary to anchor the peripheral lysosomes to the cortical actin via its effector nonmuscle myosin heavy chain IIA ^47^. Our results showed no significant changes in Rab7a mRNA and protein levels in ChA-treated cells (Supplementary Figure VIII A and B and Figure 6A). However, an increase in Arl8b and Rab3a mRNA and protein expression levels were observed in ChA-treated macrophages (Supplementary Figure VIII A and B and Figure 6 B-C). Therefore, we addressed the cellular distribution of Rab7a, Arl8b and Rab3a in peripheral enlarged LAMP-2-positive structures in single cells (Figure 6D-E). Cells treated with ChA did not present an increase in the number of lysosomes positive for Rab7a or increase in this GTPase per lysosome (Figure 6D, 6E, 6G and Supplementary Figure VIII C). In contrast, macrophages treated with ChA had more Arl8b and Rab3a-positive lysosomes which were also enriched in Arl8b and Rab3a when compared with lysosomes in control cells (Figure 6E-H). Finally, correlation of total area of the GTPase in each lysosome as function of the distance from nuclear membrane showed a random distribution for Rab7a while for Arl8b and Rab3a their area increased with distance from nuclear membrane (Figure 6I).

**Figure 6:**
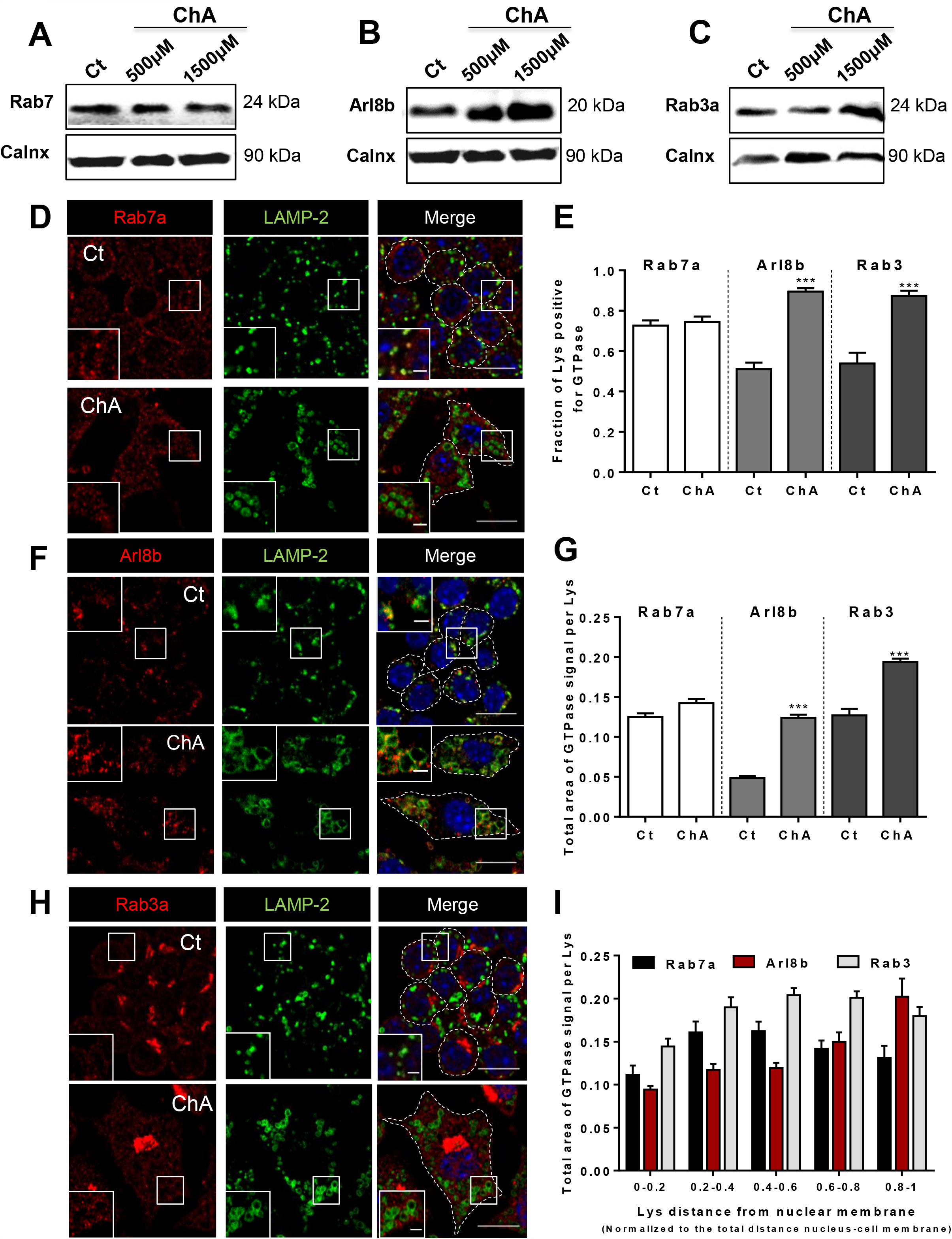
In ChA-treated macrophages the peripheral lysosomal membranes are enriched in Arl8b and Rab3a. RAW cells were incubated with 1500 µM ChA or with POPC (control cells, Ct) for 72 h. **A**-**C**. Rab7a (**A**), Arl8b (**B**) and Rab3a (**C**) protein levels in total cell lysates of control (Ct) and ChA-treated RAW cells. Calnexin was used as loading control. **D, F** and **H**. Representative confocal images of Ct and ChA-treated RAW cells co-immunostained for LAMP-2 (in green) and Rab7a (**D**, in red), Arl8b (**F**, in red) or Rab3 (**H**, in red). Nuclei were stained with DAPI (blue). Dashed lines show the edges of RAW cells. The insets are enlargements of the areas outlined with the white boxes. Scale bars, 10 μm and 2 μm in the insets. **E, G** and **I**. Quantification, using a developed ImageJ macro, of LAMP-2 vesicles positive for Rab7a, Arl8b and Rab3a (**E**) and total area of the GTPase signal per LAMP-2 vesicle (lysosomes, Lys) (**G**) in Ct and ChA-treated RAW cells. **I**. Total area of GTPases detected signal per LAMP-2 vesicle area as a function of lysosome distance from nuclear membrane. Values near 1 indicate a close proximity to the PM. Data is the mean ± SEM of three independent experients. Three independent experiments were performed and in total at least 45 cells were analysed per condition. Statistical significance was assessed by the t-test: ***, p<0.001.

### In ChA-treated cells the peripheral enlarged lysosomes are more exocytic

In recent years, the role of lysosome positioning in exocytosis, the underlying mechanisms and their physiological implications, have been largely studied ^47, 56, 60, 68, 69^. In nonsecretory cells, it was demonstrated that membrane proximal lysosomes are the organelles responsible for calcium-mediated exocytosis ^70^. Due to the peripheral lysosomal positioning in ChA-treated cells we hypothesized that these lysosomes were more exocytic than those in control cells. To assess lysosome exocytosis, we performed a flow cytometry-based assay to detect LAMP-2 at the PM ^71^. In basal conditions, without any exocytic stimulus, macrophages incubated with ChA showed higher levels of LAMP-2 at the PM than control cells (Figure 7A-C). Since effects on PM remodeling can affect LAMP-2 levels at the cell surface, ChA-treated cells were incubated at 4°C with FM4-64, a lipophilic probe that exhibits low fluorescence in water but fluoresces intensely upon binding to the outer leaflet of the PM. The cells were then shifted to 37°C to follow PM remodeling using live confocal imaging, assessed by the formation of endocytic vesicles over time. The results showed that ChA-treated macrophages did not show any delay in membrane remodeling that could contribute to the higher levels of LAMP-2 at membrane surface (Figure 7D and 7E). To further characterize the exocytic capacity of the lysosomes we evaluated their response to an exocytic stimulus such as ionomycin, a Ca^2+^ ionophore. When ChA-loaded macrophages were stimulated with ionomycin in the presence of Ca^2+^, they showed a 1.67-fold increase in LAMP-2 MFI at the cell surface in relation to stimulated control cells (Figure 7F, G). This outcome was concomitant with a significant decrease in the area and number of peripheral enlarged lysosomes (Figure 7 H-J) suggesting that enlarged lysosomes are more prone to fusion than the smaller ones. To confirm the increase in lysosome exocytosis, we also evaluated the release of the lysosomal enzyme β-hexosaminidase after addition of ionomycin and Ca^2+^. The result in Figure 7K showed a higher release of this lysosomal enzyme into the extracellular medium when cells were exposed to ChA in comparison to control cells. Altogether, these results suggested that the peripheral lysosomes in macrophages treated with ChA were more exocytic.

**Figure 7:**
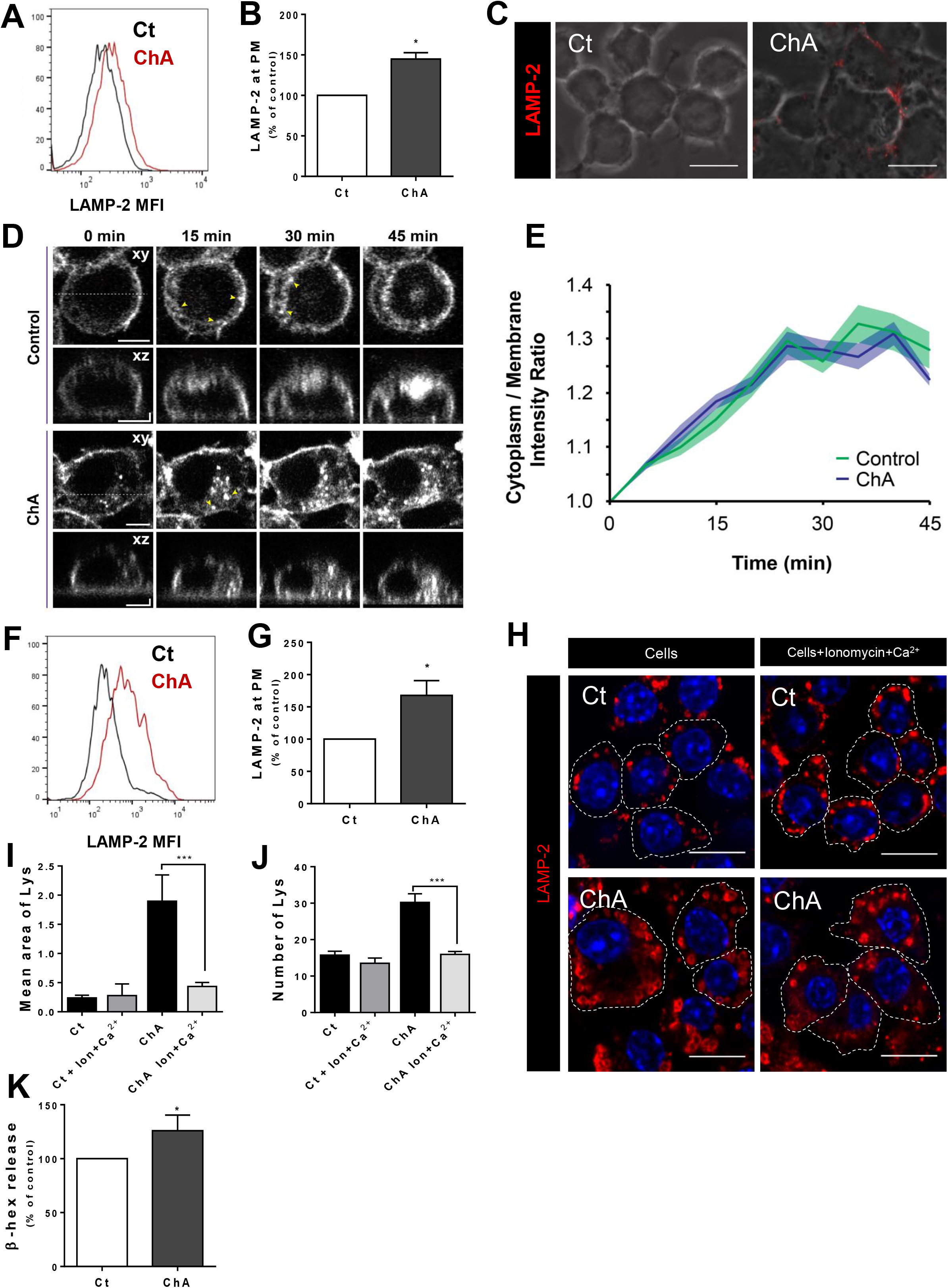
In presence of ChA the peripheral dysfunctional lysosomes are exocytic. RAW cells were incubated with 1500 µM ChA or with POPC (control cells, Ct) for 72 h. **A**. Representative histogram of LAMP-2 levels at the cell surface of Ct and ChA-treated macrophages. The values in the x-axis indicate the LAMP-2 mean fluorescence intensity (MFI) of the selected cell population. **B**. LAMP-2 levels, quantified by Flow Cytometry, at the cell surface of non-permeabilized RAW cells treated with POPC or ChA. The PI-positive cells were excluded. **C**. Representative confocal images of control and ChA-treated cells exhibiting LAMP-2 staining at the cell surface without permeabilization. The fluorescent images are merged with the corresponding DIC images. Scale bars, 10 μm. **D**. Representative images of FM4-64 fluorescence in live cells in function of time. Scale bars xy 5 μm, yz 1 μm. Arrowheads point to FM4-64 internalized vesicles and the dashed lines in the xy projections indicate the orthogonal sections shown in xz projections. **E**. Ratio of cytoplasm/plasma membrane (PM) FM4-64 fluorescence. FM4-64 fluorescence, at 15 min incubation of cells on ice, was normalized to 1 (0 min chase time). Results are mean ± SEM of three independent experiments. SEM is represented by the color area around the mean line (in total at least 45 cells were analysed). **F**. Histogram showing LAMP-2 levels at membrane surface of RAW cells stimulated with ionomycin in presence of Ca^2+^. **G**. Quantification of LAMP-2 at the PM. *, p<0.05 estimated by a t-test. **H**. Representative images of control and ChA-treated macrophages immunostained for LAMP-2 (red) before and after ionomycin treatment in presence of Ca^2+^. Nuclei were stained with DAPI (blue). Dashed lines show the edges of RAW cells. Scale bars, 10 μm. **I and J**. Quantification of the area (**I**) and number (**J**) of LAMP-2-vesicles per cell before and after ionomycin treatment in presence of Ca^2+^ (at least 45 cells were analysed by condition). ***, p<0.001 calculated using one way ANOVA-Kruskal-Wallis test. **K**. Graph showing percent of β-hexosaminidase (β-hex) release upon ionomycin treatment in presence of Ca^2+^. Data are the mean ± SEM of three independent experiments. Statistical significance was assessed by the t-test: *, p<0.05.

Some of the cellular machinery associated with lysosome exocytosis has already been identified ^47, 72^. We, therefore, investigated if lysosomes in ChA-treated cells mirror the signature of the already described secretory lysosomes. We started by addressing the expression of CD63, a tetraspanin of secretory lysosomes also known as LAMP3, required for the recruitment of the Ca^2+^ sensor synaptotagmin VII to lysosomes ^73^, by western-blot. The total CD63 levels were higher in ChA-treated macrophages (Supplementary Figure IX A and C). In single cell analysis the fraction of LAMP-2 vesicles positive for CD63 as well as the total signal of the latter per lysosome was also higher in ChA-treated than in control cells (Supplementary Figure IX D and F-H). Interestingly, ChA also induced an increase in CD63 accumulation at the PM (Supplementray Figure IX D). Then, we investigated the protein levels and the intracellular distribution of the endogenous (v)-SNARE protein, VAMP7 (also known as Ti-VAMP). This protein localizes in late endosomal compartments and is involved in the exocytosis of non- and secretory lysosomes in various cell types ^74-76^. As shown in Supplementary Figure IX B and C, ChA-treated cells presented higher VAMP7 expression levels when compared with control macrophages. These results are well correlated with the increase of lysosomes positive to VAMP7 (Supplementary Figure IX F), the increased number of VAMP7 foci, and total signal area in the lysosomal membranes from ChA-treated macrophages (Supplementary Figure IX G-H). Finally, by correlating the levels of CD63 and VAMP7 per lysosome and the distance of these organelles to the PM, we observed an increase of both secretory proteins in the more peripheral LAMP-2 vesicles (Supplementary Figure IX I). Thus, the increase in VAMP7 and CD63 levels of peripheral lysosomes induced by ChA can promote lysosome fusion with the PM and the subsequent release of their luminal content.

### MARS-Sequence Analysis confirms that ChA is pro-atherogenic and affects lysosomal physiology

Finally, to prove that ChA was pro-atherogenic and was affecting the lysosome pathway we performed MARS-seq analysis of macrophages previously exposed to the lipid. MARS-seq analysis was performed on macrophages that were incubated with ChA:POPC liposomes for 72 h. Clustering analysis of gene expression from differentially expressed genes (FDR < 0.05, and absolute fold change difference significantly higher than 1.1) showed that ChA-treated cells presented a different transcriptome profile when compared to control cells (Figure 8A). In order to identify the pathways significantly enriched for genes that are differentially expressed in ChA-treated cells, we performed a gene set enrichment analysis using the signalling and metabolic pathways available for mouse in KEGG database ^44^. Supplementary Table VI shows macrophage-related pathways affected by ChA. Interestingly, from the 99 genes of the lysosome pathway that were analysed, 11 genes were significantly differentially expressed with a *p*-value <0.05. Among them cathepsins, cholesterol exporters, components of the V-ATPases and CD63 (Figure 8B) were upregulated following ChA treatment.

**Figure 8:**
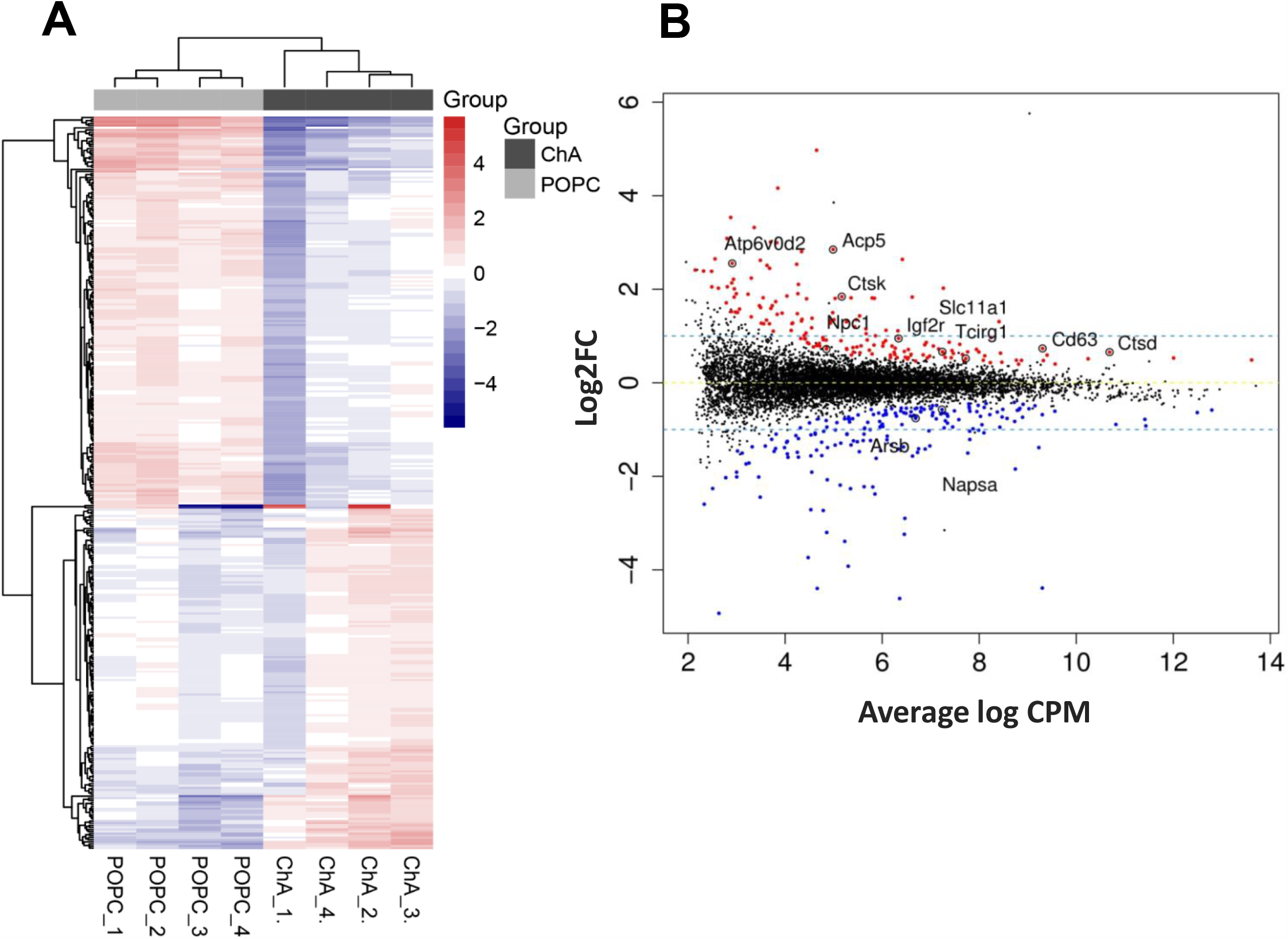
ChA drives a different gene profile in macrophages. RAW cells were incubated with 1500 µM ChA or with POPC (control cells, Ct) for 72 h. **A**. Global analysis of the differential expressed (DE) genes between macrophages treated with ChA or POPC (control cells). **B**. Mean-Difference Plot of Expression Data. Each point represents one gene plotted as log2 CPM (counts per million - measure of expression level) versus log2FC (fold change) values. Red and blue indicate DE genes with False Discovery Rate (FDR) < 0.05 and absolute fold change difference significantly higher than 1.1. DE genes from the lysosomal pathway are highlighted.

## DISCUSSION

Macrophages are mainly responsible for the phagocytic removal of LDL particles retained and subsequently oxidized in the arterial subendothelial *intima*. However, in the course of this process, a progressive lysosome dysfunction leads to an inefficient digestion of the Ox-LDL in lysosomes, causing lipidosis, subsequent cell apoptosis, and defective cellular debris clearance (reviewed in ^77^). Understanding the chemical etiology of this macrophage dysfunction in atherosclerotic plaques would be important in possible prophylactic approaches or reversal of atheroma formation.

Among the several risk factors already identified, the blood LDL-cholesterol level is one of the main indicators of CVD risk. However, cholesterol-LDL or any of the existing risk factors of atherosclerotic CVD, are good predictors for unstable atherosclerotic plaques. Cholesteryl esters are also one of the main components of LDL particles and, due to the polyunsaturation of their fatty acid component, are susceptible to oxidation. Our lipidomic results show that linoleic acid constitutes 50 % of the total, and 77 % of the polyunsaturated fatty acid components of cholesteryl esters in plasma (unpublished results). Thus, we estimated that ChA would be one of the main end-products of cholesteryl linoleate oxidation increased in CVD. In the present work we were able to show that ChA is the most prevalent ChE in human plasma. As far as we know, this is the first time that these lipids have been detected and reported to be increased in the lipidome of CVD patients. As discussed earlier, the detected concentration of about 1.5 µM ChA in the plasma of CVD patients implies a localized concentration ≥ 0.1 M ChA in the polar lipidic surface of the atheroma which the macrophages unsuccessfully attempt to clear up. There is a vast amount of literature on Ox-PL, oxidized cholesterol, and oxidized cholesteryl ester hydroperoxides (Ox-CE) and their involvement in atherosclerosis ^21, 78-80^, but information on ChE in the pathology of artherosclerosis is lacking.

We had previously demonstrated that ChS, a commercially available ChE, was able to induce cell lipidosis and lysosome dysfunction in macrophages ^36, 37^. Here, we show that the most prevalent ChE found in human plasma, ChA, also causes lysosome dysfunction, lipidosis, and exocytosis of these dysfunctional organelles. Since the lipid and other cargo in the dysfunctional exocytosed lysosomes is essentially undigested, its deposition in the arterial subendothelial intima media could be part of the initiation process of atheroma formation but also, more importantly, a self-perpetuate the pathological process.

The inflammatory profile of oxidized lipids has been intensively studied in the literature ^79, 80^ and taking into account our transcriptomic profiling data, ChA also seems to be inflammatory (in preparation). Nevertheless, the novelty of this lipid is its strong impact on the lysosome pathway causing the overexpression of many lysosome-related genes. The transcriptomic data is well correlated with lysosome phenotypic changes. Indeed, ChA induces striking changes in lysosome positioning and morphology having impact in their function and inducing lysosome biogenesis and exocytosis. The peripheral localization of lysosomes in ChA-treated macrophages was unexpected because lipidotic macrophages share similarities with those found in some lysosomal storage diseases (LSD). In LSD, these acidic organelles are normally localized in the perinuclear region of the cells. For example, in Niemann Pick C1, a LSD associated with mutations in NPC1 (a cholesterol exporter), there is an accumulation of FC in lysosomes and these organelles become mainly perinuclear ^81^. Furthermore, in lysosomes with high content of cholesterol ^82^, the cholesterol-interacting domain of oxysterol-binding protein-related protein 1 long (ORP1L), ORD, aquires a different conformation in the lysosomal membrane and Rab7a-effector RILP recruits the dynein-dynactin motor complex to facilitate minus-end transport ^66, 67, 83^. ChA and cholesterol have some structural similarities: both are amphiphilic molecules and partition via passive diffusion and trans-membrane translocation into all cell membranes. Thus, as cholesterol, ChA is expected to affect membrane lateral packing density. Additionally, ChA has a polar head-group that acquires a negative charge at neutral pH and, thereby, alters the membrane surface charge. These alterations in the biophysical properties of ChA-inserted membranes will affect the association of proteins with these modified membranes and could explain, at least in part, the low density of Rab7a and the high density of Arl8b in the lysosomal membranes, which leads to the peripheral lysosomal localization ^56^. In addition, the low density of Rab7 in the enlarged lysosomes can also be related to the increase in lysosome luminal pH. Rab7a-effector RILP is responsible for the recruitment and stability of the V1G1, a component of the lysosomal V-ATPase ^84^, important for the V-ATPase activity, the proton transporter responsible for lysosomal acidification. Changes in the lysosomal luminal pH can also be expected to affect activity of acid hydrolases as well as membrane trafficking and cargo sorting ^85-87^. It is, therefore, expected that transport of cargo that is targeted to lysosomes and its degradation, as well as cargo sorting at the early stages of the endocytic pathway are affected upon exposure of cells to ChA. The decrease in the hydrolytic capacity of lysosomal enzymes followed by cargo accumulation can explain lysosomal enlargement. As described by other groups in different experimental settings ^58^, lysosomal biogenesis can also contribute to lysosome swelling. In our case, considering the increase in lysosome number we do not believe that lysosome biosynthesis contributes to lysosome enlargement. However, it is possible that ChA affects the balance in lysosome fusion-fission cycling, contributing also to lysosome enlargement.

It is known that only a small fraction of the lysosomes at the periphery of non-secretory cells are exocytic ^70^. One of the main novelties of the present work is that the peripheral enlarged dysfunctional lysosomes are more prone to fuse with the PM. This feature can be explained by the enrichment of lysosomal membranes in VAMP7 that mediates lysosome fusion at the PM and CD63, a tetraspanin of secretory lysosomes, that also supports the exocytic properties of these organelles. This aspect was never reported in the pathology of atherosclerosis. Thus, it seems reasonable to envision that lysosome exocytosis and *de novo* biogenesis of these organelles are the cellular responses to cope with aberrant lysosomes upon ChA treatment. However, while lysosome biogenesis can ameliorate the pathology, exocytosis of the aberrant lysosomes can have the opposite effect. Indeed, lysosome exocytosis can create an acidic local hub, secreting cathepsins among other enzymes that contribute to the development and progression of atherosclerosis by influencing extracellular matrix turnover, inflammation and apoptosis ^88-90^. Furthermore, the secretion of undigested cargo including ChA can contribute to increased “danger (or damage) associated molecular patterns” (DAMPs) exposure triggering a variety of immune responses and expression of proinflammatory genes as have been described for Ox-PL and Ox-CE ^79, 80^.

Considering that only a few reports have linked macrophage dysfunction in atheroma with the final component of the cellular traffic system, the lysosome, the data reported here suggests that interventions aiming at restoring lysosomal function may be useful in atherosclerosis treatment.

## ACKNOWLEDGEMENTS

We thank the nursing staff at the Hospital Santa Cruz, in particular Edite Tomás Mateus and Manuel Belo Costa for their enthusiastic involvement in this project and help in blood collection. We acknowledge the UC-NMR facility for obtaining the NMR data (www.nmrccc.uc.pt) and the technical support of the CEDOC Microscopy and Flow cytometry facilities. We also would like to thank the Life Science Core Facilities at the Weizmann Institute of Science. Finally, we thank Liliana Alves for the cell culture support and André Marques and Jorge Silva for the careful reading and improvement of this manuscript.

## Non-standard Abbreviations and Acronyms

ACS: Acute coronary syndrome
ChA: Cholesteryl hemiazelate
ChE: Cholesteryl hemiesters
ChS: Cholesteryl hemisuccinate
SAP: Stable angina pectoris

