## Supplementary Material for "Cholesteryl Hemiazelate Induces Lysosome Dysfunction and Exocytosis in Macrophages"

**Authors Affiliations:**[1] CEDOC, NOVA Medical School | Faculdade de Ciências Médicas, Universidade NOVA de Lisboa, 1169-056 Lisboa, Portugal. [2] CQC and Department of Chemistry, University of Coimbra, 3004-535 Coimbra, Portugal. [3] Department of Cell Biology, UCL Institute of Ophthalmology, London, U.K [4] Department of Biomolecular Sciences, Weizmann Institute of Science, Rehovot, Israel. [5] Lipotype GmbH, Tatzberg 47, 01307 Dresden, Germany. [6] Hospital Santa Cruz, Centro Hospitalar de Lisboa Ocidental, Av. Prof. Dr. Reinaldo dos Santos, 2790-134 Carnaxide, Portugal. [7] Department of Biomedical Science & Centre for Membrane Interactions and Dynamics. University of Sheffield, UK. [#] Current address: UCIBIO-REQUIMTE, Departamento de Ciências da Vida, Faculdade de Ciências e Tecnologia, Universidade Nova de Lisboa, 2829-516 Caparica, Portugal. [§] Current address: Centre de Recherche, Institut Curie, 26 rue d'Ulm, 75248 Paris Cedex 05, France

#### SUPPLEMENTARY METHODS

##### Cholesteryl hemiesters synthesis

**General Remarks:** Thin-layer chromatography (TLC) analyses were performed using precoated silica gel plates. Flash column chromatography was performed with silica gel 60 as the stationary phase. $^{1}H$ NMR spectra were recorded on an instrument operating at 400 MHz and $^{13}C$ NMR spectra were recorded on an instrument operating at 100 MHz. The spectra were recorded in $CDCl_3$ as solvent, chemical shifts are expressed in parts per million (ppm) relatively to internal tetramethylsilane (TMS), and coupling constants (*J*) are in hertz (Hz). Infrared spectra (IR) were recorded in a Fourier Transform spectrometer coupled with a diamond Attenuated Total Reflectance (ATR) sampling accessory. High-resolution mass spectra (HRMS) were obtained on an electrospray (ESI) APEX-Qe or microTOF mass spectrometer. Melting points were determined in open glass capillaries. Azelaic acid, acetyl chloride, glutaric anhydride, cholesterol and ammonia were purchased from commercial sources and used as received. Pyridine was dried over KOH. Ethanol and methanol were purified by distillation. Chloroform was distilled and passed through a column of basic alumina.
Cholesteryl hemiesters were prepared following the synthetic pathway outlined in Supplementary Figure I, by using an adapted procedure, described for the synthesis of cholesteryl hemisuccinate[1].

**Cholesteryl hemiazelate (ChA, cholesteryl *O*-(8-carboxyoctanoyl)):** Azelaic anhydride was prepared by the reaction of azelaic acid with acetyl chloride as described in the literature[2]. A solution of cholesterol (2.62 g, 6.78 mmol) and azelaic anhydride (3.00 g, 17.63 mmol) in dry pyridine (27 mL) was heated under reflux for 7 h. Pyridine was removed under reduced pressure and the crude product was purified by flash chromatography [chloroform/methanol/ammonia (50:5:0.25, v/v)] giving cholesteryl hemiazelate as a fluffy solid. Further recrystallization from ethanol gave cholesteryl hemiazelate as a white solid (2.16 g, 57%). m.p. 79.5-80.4 ºC. IR (ATR) *v* 1168, 1234, 1466, 1702, 1730, 2850, 2866 and 2932 cm$^{-1}$. RMN $^{1}H$ ($CDCl_3$, 400 MHz) $\delta$ = 0.68 (s, 3H), 0.86 (d, *J* = 1.7 Hz, 3H), 0.87 (d, *J* = 1.7 Hz, 3H), 0.91 (d, *J* = 6.5 Hz, 3H), 0.94-1.65 (m, 35H), 1.81-1.87 (m, 3H), 1.93-2.02 (m, 2H), 2.25-2.36 (m, 6H), 4.58-4.65 (m, 1H), 5.37-5.38 (m, 1H). RMN $^{13}C$ ($CDCl_3$, 100 MHz) $\delta$ = 11.9, 18.7, 19.3, 21.0, 22.6, 22.8, 23.8, 24.3, 24.6, 24.9, 27.8, 28.0, 28.2, 28.9, 31.9, 31.9, 33.8, 34.6, 35.8, 36.2, 36.6, 37.0, 38.1, 39.5, 39.7, 42.3, 50.0, 56.1, 56.7, 73.7, 122.6, 139.7, 173.3, 179.1. HRMS (ESI) *m/z*: Calcd for $C_{36}H_{61}O_4$ $[M+H]^+$ 557.45644; found 557.45615.

**Cholesteryl hemiglutarate (ChG, cholesteryl *O*-( 4-carboxybutanoyl)):** A solution of cholesterol (3.71 g, 9.60 mmol) and glutaric anhydride (1.90 g, 16.32 mmol) in dry pyridine (35 mL) was heated under reflux for 6 h. Pyridine was removed under reduced pressure and the crude solid product was triturated with methanol. The filtered solid was further purified by flash chromatography [chloroform/methanol/ammonia (50:5:0.25, v/v)] giving cholesteryl hemiglutarate as a white solid (1.79 g, 37%). m.p. 124.4-125.9 ºC. IR (ATR) *v* 1174, 1282, 1376, 1428, 1702, 1729, 2866 and 2932 cm$^{-1}$. RMN $^{1}H$ ($CDCl_3$, 400 MHz) $\delta$ = 0.68 (s, 3H), 0.86 (d, *J* = 1.6 Hz, 3H), 0.87 (d, *J* = 1.6 Hz, 3H), 0.91 (d, *J* = 6.5 Hz, 3H), 0.94-1.57 (m, 25H), 1.80-1.88 (m, 3H), 1.92-2.05 (m, 4H), 2.30-2.45 (m, 6H), 4.58-4.66 (m, 1H), 5.37-5.38 (m, 1H). RMN $^{13}C$ ($CDCl_3$, 100 MHz) $\delta$ = 11.9, 18.7, 19.3, 19.9, 21.0, 22.6, 22.8, 23.8, 24.3, 27.8, 28.0, 28.2, 31.9, 31.9, 32.8, 33.5, 35.8, 36.2, 36.6, 37.0, 38.1, 39.5, 39.7, 42.3, 50.0, 56.1, 56.7, 74.1, 122.7, 139.6, 172.3, 177.8. HRMS (ESI) *m/z*: Calcd for $C_{32}H_{53}O_4$ $[M+H]^+$ 501.39384; found 501.39160.

##### Filipin staining and binding experiments

The binding capacity of Filipin to free cholesterol (FC) and to ChA was performed using a spectral assay. Several dilutions of POPC-FC (35:65, molar ratio) and POPC-ChA (35:65, molar ratio) liposomes were prepared and were incubated overnight with a 50 µM aqueous solution of Filipin at room temperature. Absorption spectra were obtained using the plate reader Spectramax i3x from Molecular Devices by measuring the absorbance ratio (Abs320/Abs356) and the corresponding Filipin-free lipid suspensions were used as blanks. Filipin-sterol association curves used the Abs320/Abs356 as a function of sterol concentration (FC or ChA) and were fitted with theoretical curves using the Hill Equation.

**BSA and dextran uptake and BSA trafficking**

For the endocytosis assays, cells were incubated with 400 µg/ml BSA-Texas Red or FITC-dextran for 30 or 60 min in medium without serum after lipid treatment. In order to avoid BSA degradation, $NH_4Cl$ was added to neutralize the lysosome pH 20 min after initiation of the incubation with cargo. After incubation, RAW cells were washed with PBS, detached using FACS buffer and fixed with PFA. Cargo internalization was evaluated by flow cytometry. For BSA-Texas red and dextran uptake measurements, the controls and the fluorescent-labelled samples were run on a BD FACScantoTMII flow cytometer (Becton Dickinson) cell sorter equipped with a 561nm (50 mW solid state) and 488 nm laser laser (20 mW solid state) used for Texas Red and FITC-dextran excitation, respectively. Texas Red was measured using a 610/20 nm bandpass emission filter and FITC-dextran with 530/30 nm bandpass.

To follow the intracellular transport of BSA to lysosomes, cells were incubated with 500 µg/mL BSA-Texas Red for 30 min at 37°C (pulse), then washed to remove the non-internalized BSA, and incubated again at 37°C for different time points (chase). After fixation cells were co-immunostained for LAMP-2 and EEA-1. Samples were then observed under a confocal microscope. The co-localization values were calculated using confocal single slices and ImageJ software (https://imagej.net/Citing).

**Transferrin uptake and recycling**

After lipid treatment for 72 h, cells were pulsed with 0.2 mg/mL TRITC-transferrin (Invitrogen) for 10 min, washed with PBS and chased. Then, cells were fixed after 0, 5, 10, 20 and 40 min of chase time. For the fluorescent representative images, the coverslips were mounted and imaged using confocal microscopy. To quantify transferrin uptake, the cells were detached using FACS buffer and analyzed using FACAriaTMIII (Becton Dickinson) cell sorter equipped with a 50 mW solid state 561nm laser. TRITC-transferrin was measured using a 610/20nm bandpass emission filter. Data was analysed by FlowJo software.

**Autophagic flux assays and p62 levels**

Autophagic flux was accessed by western-blot analysis measuring LC3 II turnover in the presence and absence of Bafilomycin (Baf) A1. For that purpose, RAW cells were seeded and treated with lipid during 24 or 72 h. 2 h prior to harvesting, cells were treated with 100 nM Baf A1. Whole cell lysates were run on western blots and autophagic flux was quantified by obtaining the ratio [(LC3-II:LC3-I) BAF-treated]/[(LC3-II:LC3-I) untreated]. The antibodies used were the LC-3b and p62.

#### SUPPLEMENTARY RESULTS

##### ChA affects vesicular trafficking

Differential lysosome morphology has functional implications and can reflect in an effect on cellular normal trafficking. To verify if this was indeed the case in ChA-treated macrophages, we started by challenging cells with different endocytic cargos and then followed their intracellular transport. RAW cells exposed to ChA for 72 h were loaded with BSA-Texas red, fluorescent-labelled dextran (a fluid phase marker) and rhodamine-transferrin. As shown in Supplementary Figure VI A, C and E (first column) and quantified in Supplementary Figure VI B, D and F, uptake of BSA, dextran and transferrin were not reduced by ChA treatment. These results suggested that the uptake of endocytic cargo by macrophages, whether receptor-mediated or not, was not affected by ChA. Regardless of the internalization mode, endocytosed cargo is delivered to early endosomes, where cargo sorting occurs. Cargo-specific sorting leads to distinct subsequent cargo itineraries. Cargo can be routed from the early endosomes to late endosomes and lysosomes for degradation or can be recycled back to the plasma membrane (PM). The majority of BSA and dextran is delivered to lysosomes whilst transferrin exclusively follows the recycling pathway. Then, our next step was to verify whether the recycling (transferrin) and transport of endocytic cargo (BSA-Texas Red) to lysosomes were affected in RAW cells exposed to ChA. To measure the rate of transferrin recycling, ChA-loaded RAW cells were chased, after a 10 min pulse, for different time points as indicated in graph abscissa (Supplementary Figure VI G). As shown in Supplementary Figure VI E and G, at all chase times transferrin fluorescence in ChA-treated RAW cells was higher than in control cells, reflecting a delay in transferrin-receptor recycling. Moreover, delays in early recycling are expected to have a negative impact in the endocytic pathway and as can be observed in Supplementary Figure VI H-I transport of BSA to the lysosomes was slightly delayed. Cargo delivery to lysosomes was quantified in fixed cells immunostained for Early Endosome Antigen 1 (EEA1, a marker for early endosomes) and LAMP-2 after challenging the cells for 30 min with BSA followed by different chase times as indicated (Supplementary Figure VI I). An increase in the colocalization of internalized cargo and EEA-1 and a decrease in LAMP-2-acquisition were indications of a delay on cargo delivery to lysosomes. The delay on cargo transport in the degradative pathway was more pronounced at 60 min chase where in control cells 97 ± 0.8% of the BSA positive vesicles were positive for LAMP-2 versus 89 ± 1.7 % for ChA-teated cells. Altogether, our data suggested that changes in lysosome morphology affects vesicular transport at early stages of the process since recycling of early/sorting- endosomes components was reduced. This outcome could have an impact in the endosome maturation process as described for other experimental settings[3].

#### SUPPLEMENTARY FIGURES

#### Supplementary Figure I

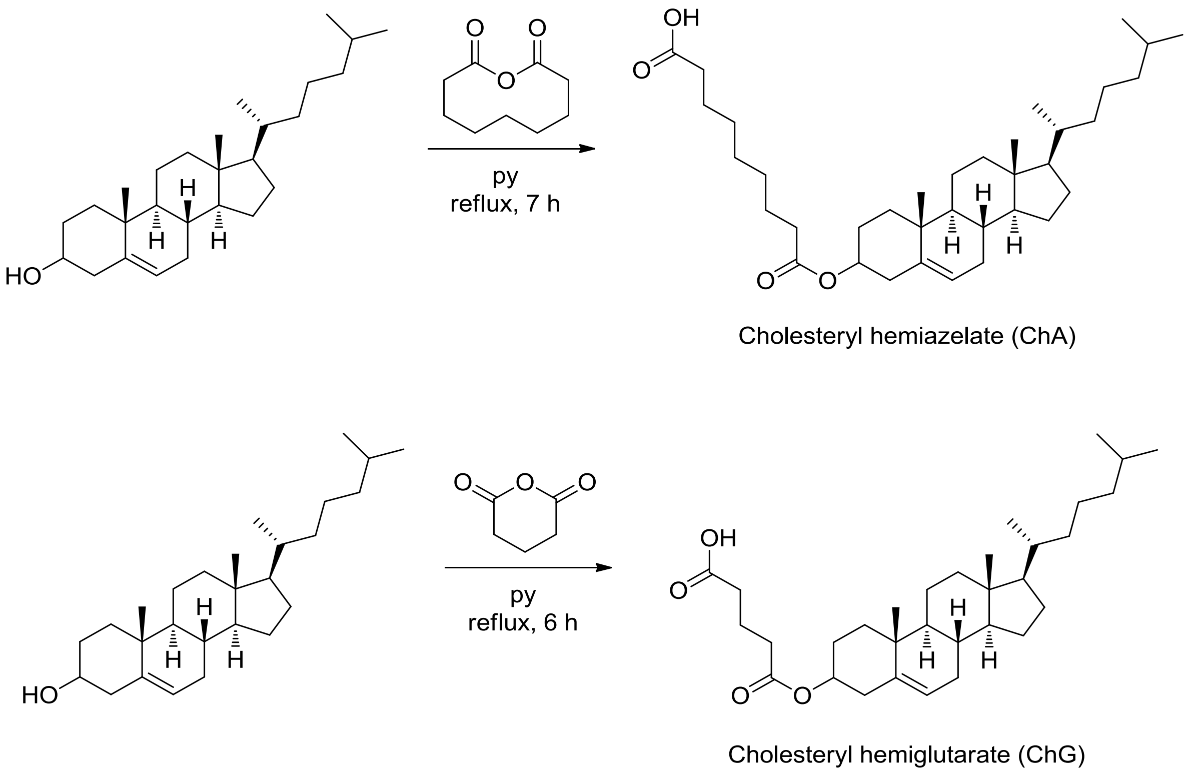


**Supplementary Figure I.** Scheme of the Synthetic strategy towards cholesteryl hemiazelate (ChA, cholesteryl O-(8-carboxyoctanoyl) and cholesteryl hemiglutarate (ChG, cholesteryl O-( 4-carboxybutanoyl) ).

#### Supplementary Figure II

A

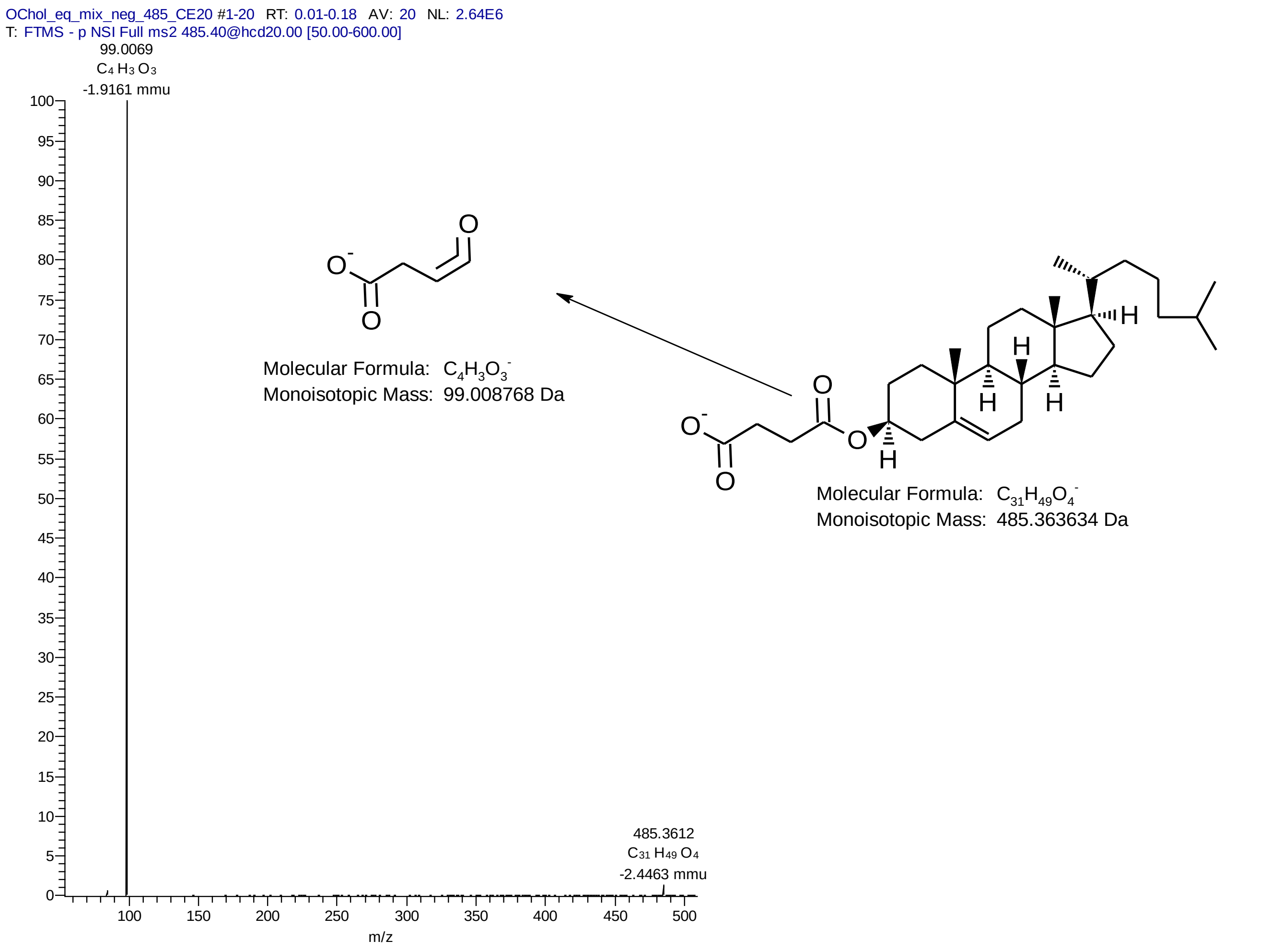


B

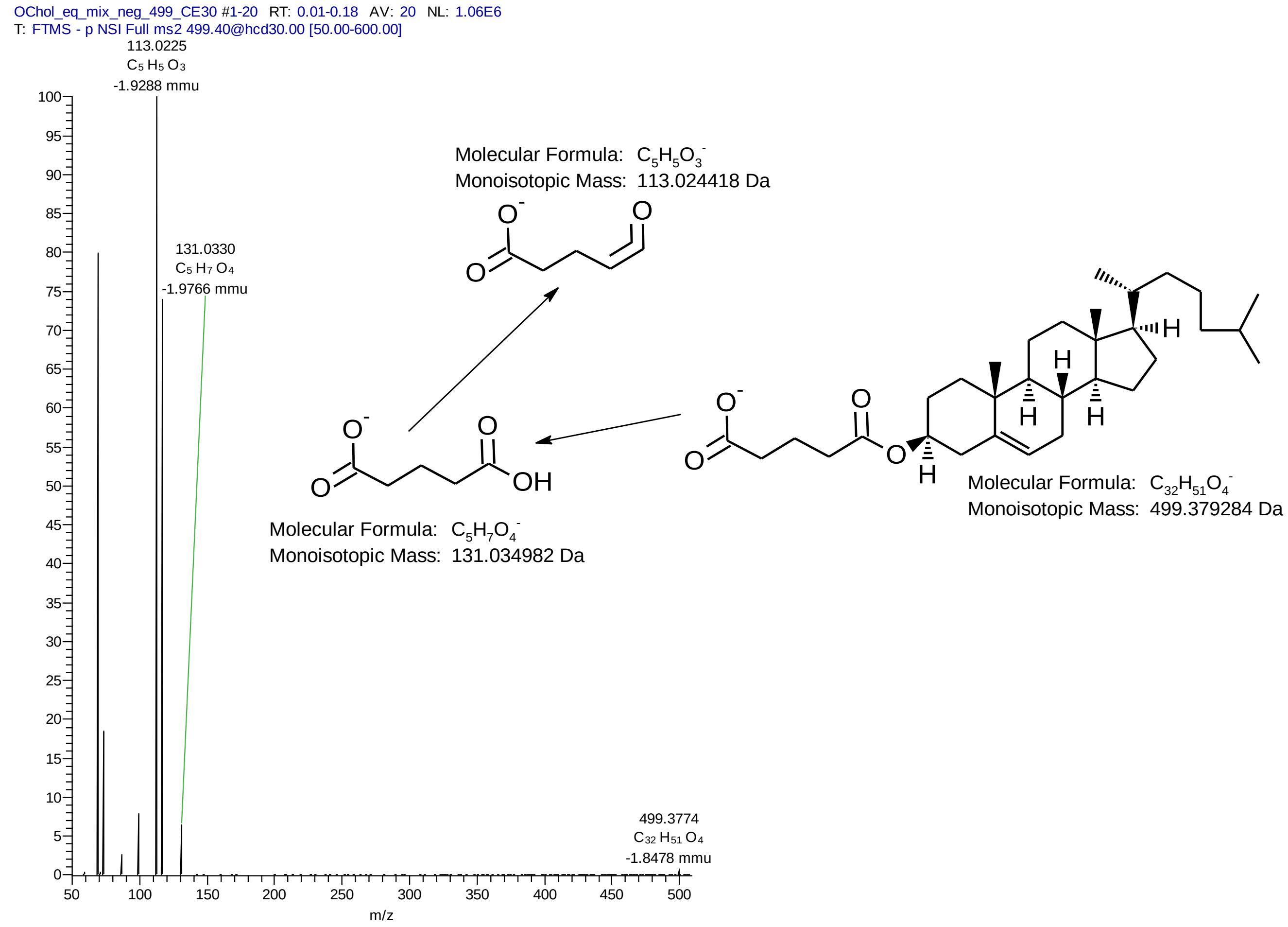

### Supplementary Figure II Cont.

**C**

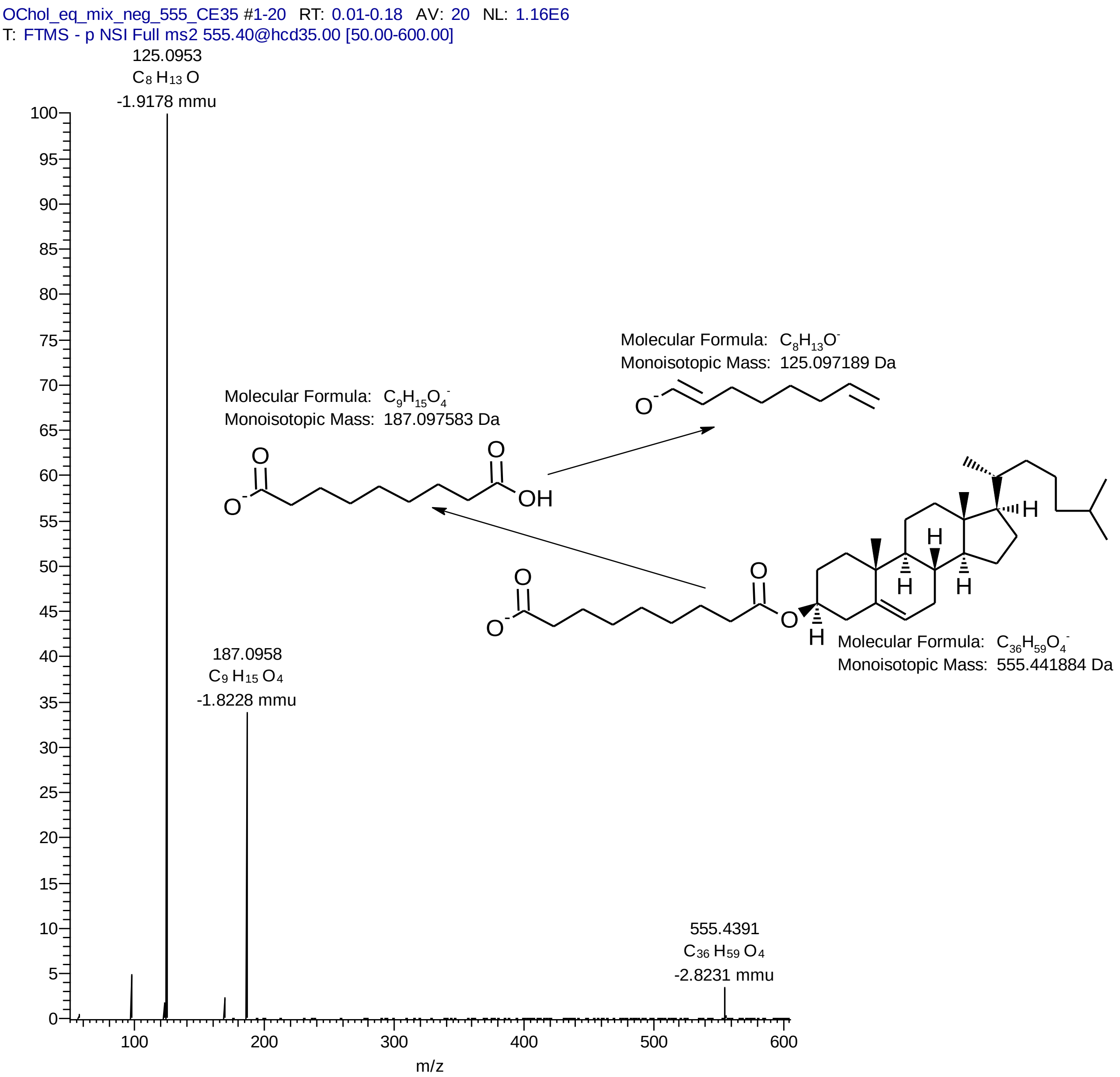


**Supplementary Figure II.**: MSMS spectra of **A.** ChS $[M-H^+]^-$ ion (at normalized ChE= 20%) and proposed fragmentation; **B.** ChS $[M-H^+]^-$ ion (at normalized CE= 30%) and proposed fragmentation, and **C.** ChA $[M-H^+]^-$ ion (at normalized ChE= 35%) and proposed fragmentation.

#### Supplementary Figure III

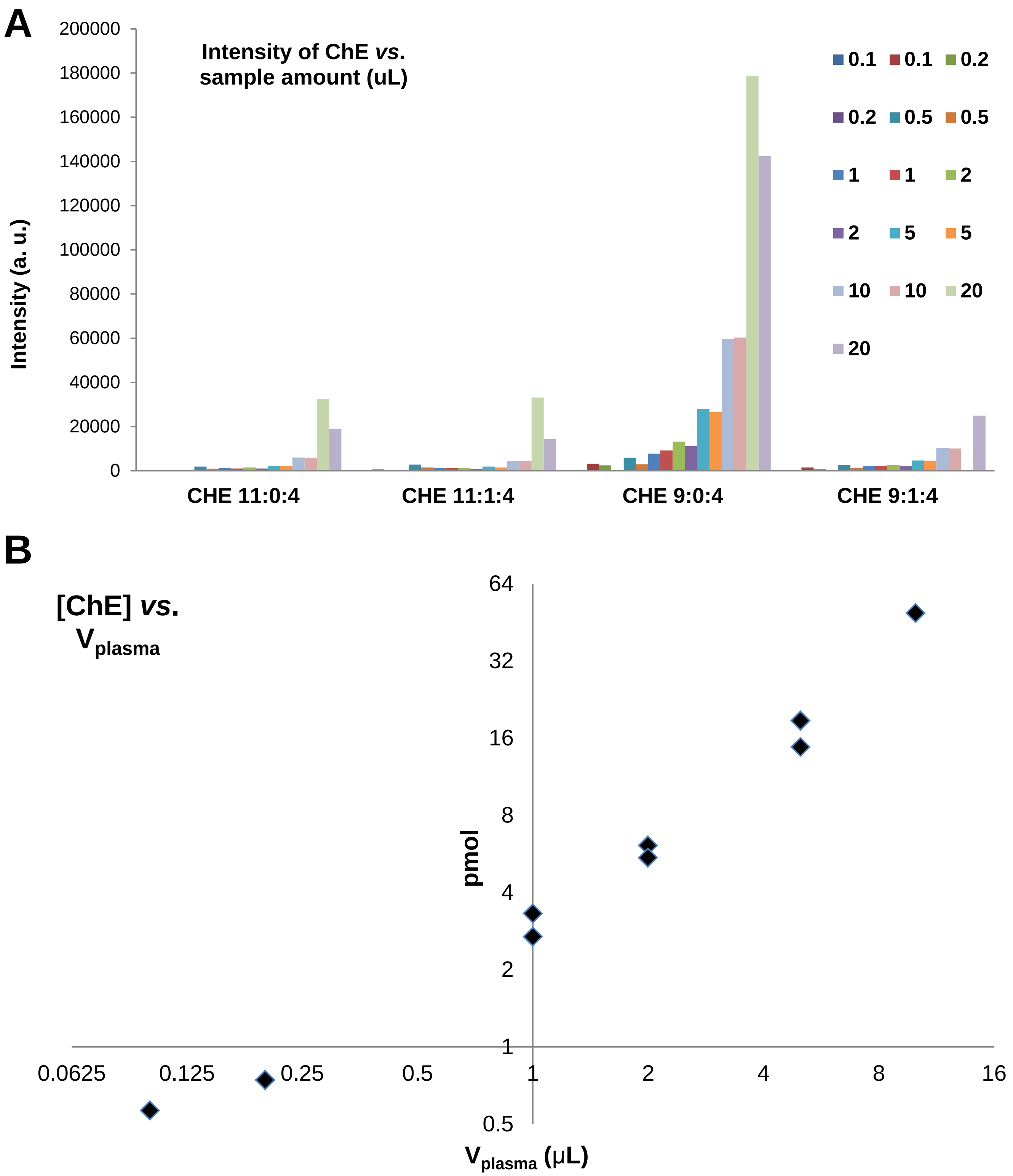


**Supplementary Figure III**. **A.** Different cholesteryl hemiesters (ChE) species observed in the test plasma sample and the dependence of their intensity with the volume of sample extracted (**B**).

#### Supplementary Figure IV

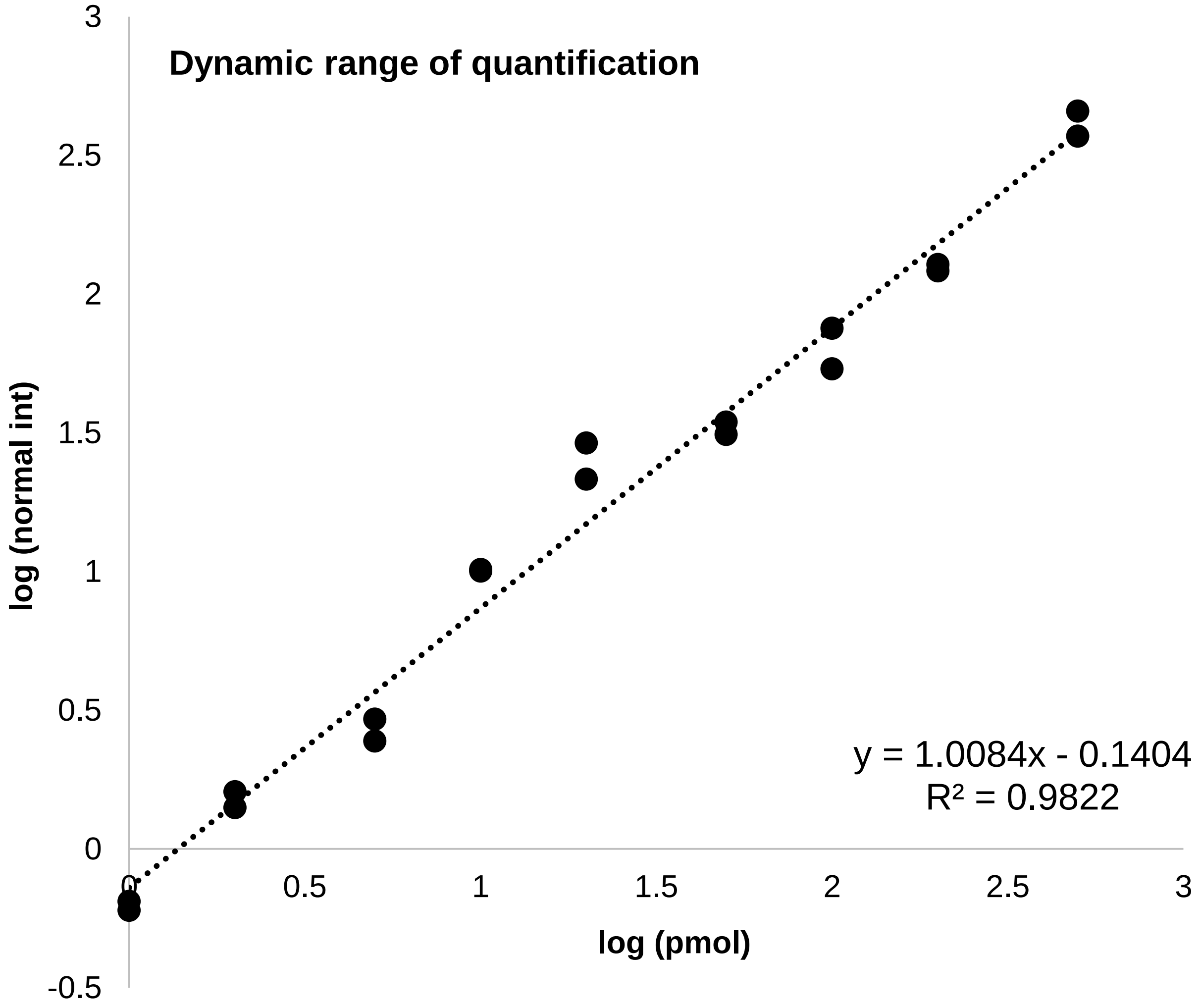


**Supplementary Figure IV.** ChE response with increasing amounts of standard added. The limit of quantification determined is 1 µM and the dynamic range is >500.

#### Supplementary Figure V

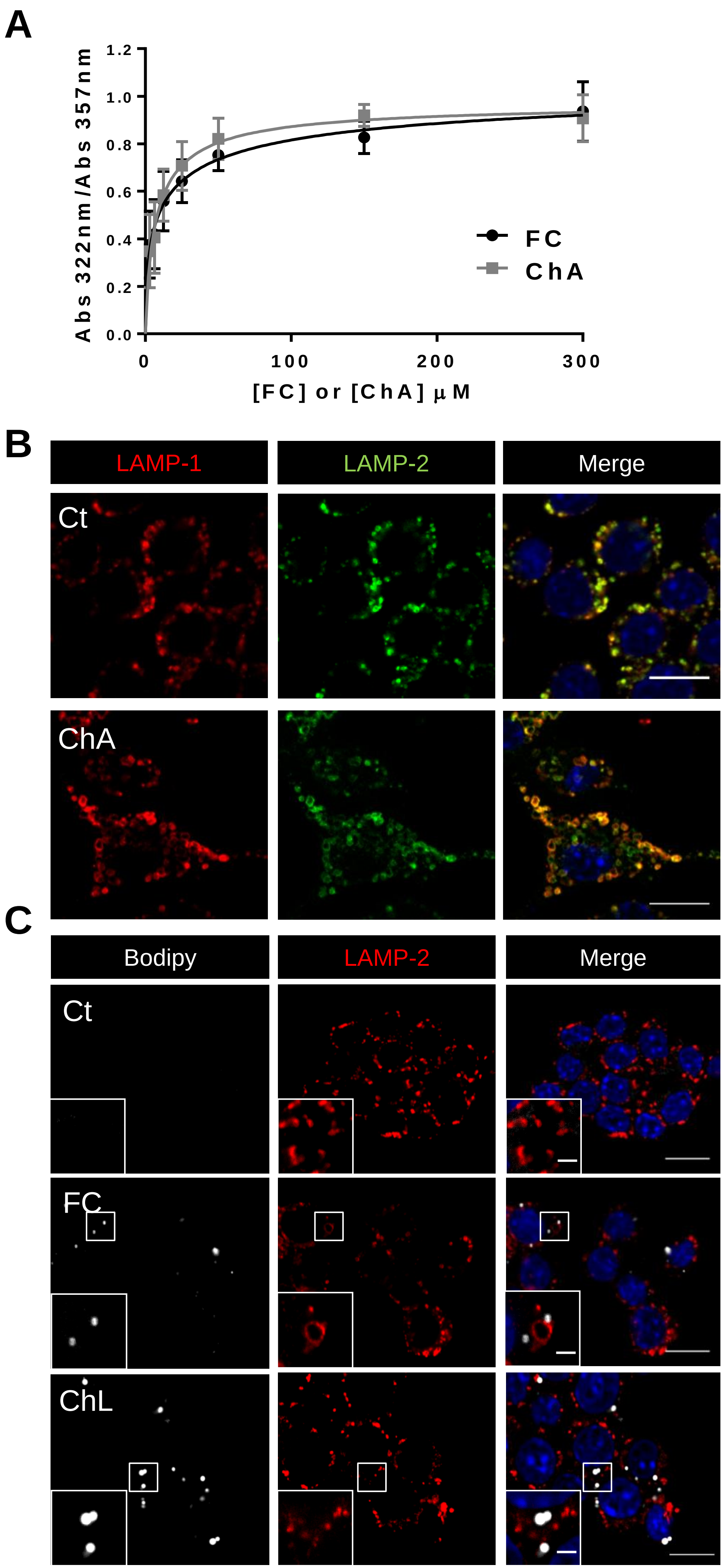

**Supplementary Figure V. Filipin binds to ChA and LAMP-1 and LAMP-2 present the same distribution in ChA-loaded macrophages. Free cholesterol and cholesterol linoleate do not induce changes in lysosome morphology.**

**A.** Filipin association curve with free cholesterol (FC), POPC-FC (35:65, molar ratio), and ChA, POPC-ChA (35:65, molar ratio). Filipin binds to both steroids with a similar binding constant. The graph shows the absorption intensity ratio ($A_{320}/A_{356}$) for Filipin as a function of FC (●, black line) or ChA (■, grey line) using Hill equation for theoretical fitting. The obtained dissociation constant is 95 μM for the Filipin-FC complex and 100 μM for the Filipin-ChA complex, with a Hill coefficient of 1.5 and 1.6, respectively. Data show the mean ± SEM of four independent experiments. **B.** Representative immuno-flourescence images of LAMP-1 (red color) and LAMP-2 (green color) distribution in RAW cells. **C.** Effect of FC and Cholesteryl linoleate (ChL) on lysosome morphology, visualized by LAMP-2 immunostaining (red color), and on neutral lipid accumulation visualized by Bodipy staining (white color). DAPI was used to visualize nuclei (blue color). RAW cells were treated for 72 h with 1500 μM of FC:POPC liposomes (65:35, molar ratio) or with 1500 μM of ChL:POPC emulsions. Scale bars, 10 μm and 2 μm in the insets.

#### Supplementary Figure VI

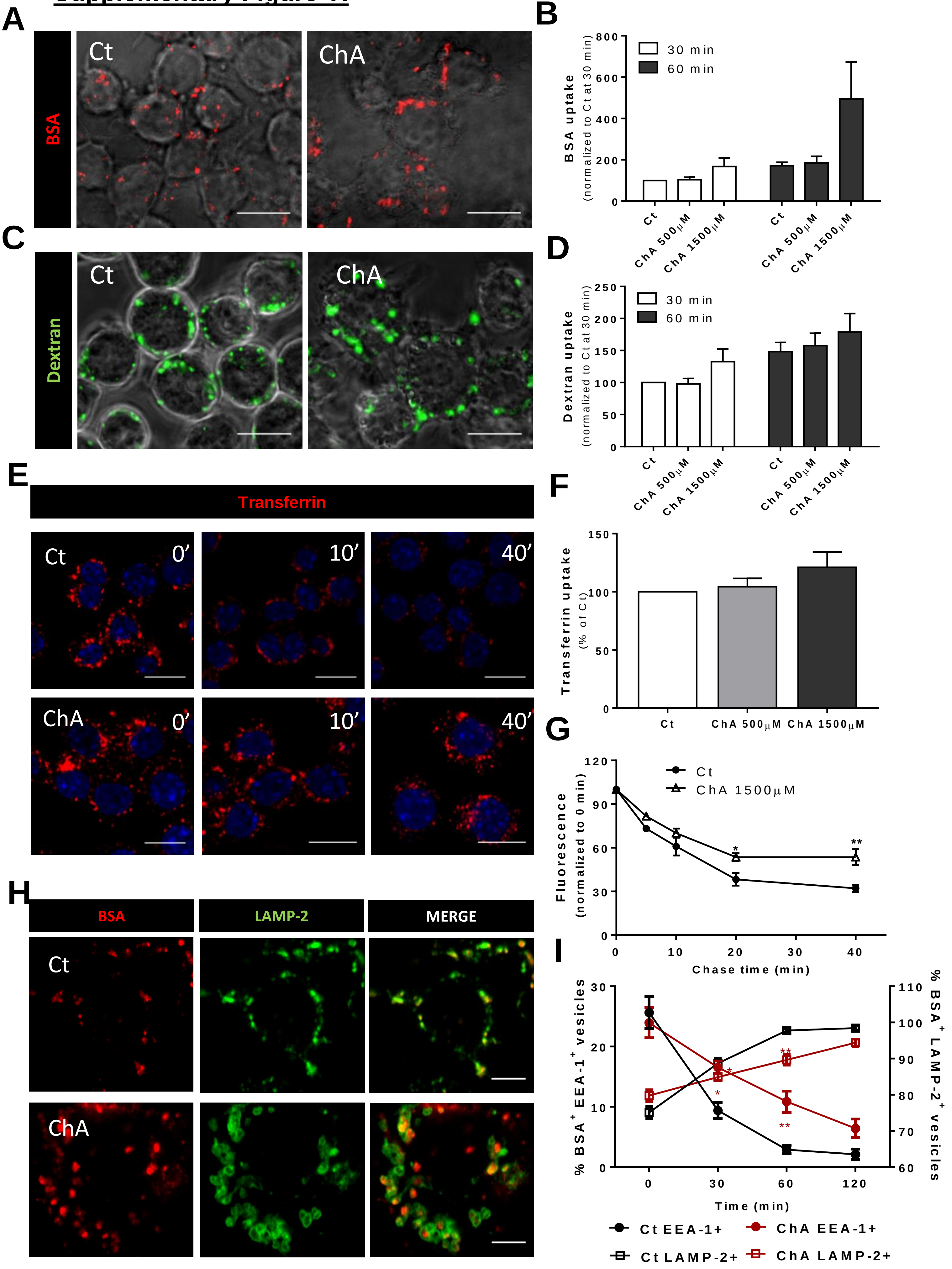

**Supplementary Figure VI. ChA affects vesicular transport in RAW cells. A.** Representative confocal images of RAW macrophages incubated with 1500 µM of ChA and with POPC liposomes (control cells, Ct) for 72 h and loaded with BSA-Texas Red for 60 min. To avoid BSA degradation the lysosomal pH was neutralized with ammonium chloride that was added 20 min after the endocytic cargo. The fluorescent images are merged with the corresponding DIC images. **B.** Quantification of the BSA-Texas Red-uptake by Flow Cytometry. Results represent the mean ± SEM of four independent experiments. **C.** Confocal images of RAW cells fixed after 60 min incubation with FITC–dextran, a fluid phase marker. The fluorescent images are merged with the corresponding DIC images. **D.** Effect of ChA on dextran internalization in RAW fixed cells assessed by Flow Cytometry. Results are the mean ± SEM of four independent experiments. **E.** Representative confocal images of Rhodamine-transferrin uptake and recycling in ChA-treated RAW cells. Transferrin recycling was assessed by chasing the cells for different time points after a 15 min pulse (0 min chase) as indicated. Nuclei were stained with DAPI (blue). **F.** Quantification of ChA on Rhodamine-transferrin uptake evaluated by Flow Citometry. **G.** Quantification of ChA on Rhodamine-transferrin recycling assessed by Flow Citometry. The values are the mean ± SEM of three independent experiments. **H.** Representative confocal images of LAMP-2 distribution in RAW cells incubated with BSA-Texas Red for 30 min and then chased for 60 min. The first and the second columns show BSA (red), and LAMP-2 (green) staining, respectively. The third column is composite of the BSA and LAMP-2. **I.** Effect of ChA on the BSA-containing vesicles trafficking assessed by the loss of EEA-1 and acquisition of LAMP-2. POPC- (black lines) and ChA- treated (red lines) RAW cells were pulsed with BSA-Texas Red during 30 min and then chased for different time points as indicated in the graph abcissa. The colocalization between BSA-containing vesicles and EEA-1 (●) or LAMP-2 (□) was quantified in confocal images by ImageJ. Data represent the mean ± SEM of three independent experiments. Statistical significance was assessed by a one-way ANOVA-Kruskal-Wallis test: *, $p < 0.05$; **, $p<0.01$. Scale bars, 10 µm.

#### Supplementary Figure VII

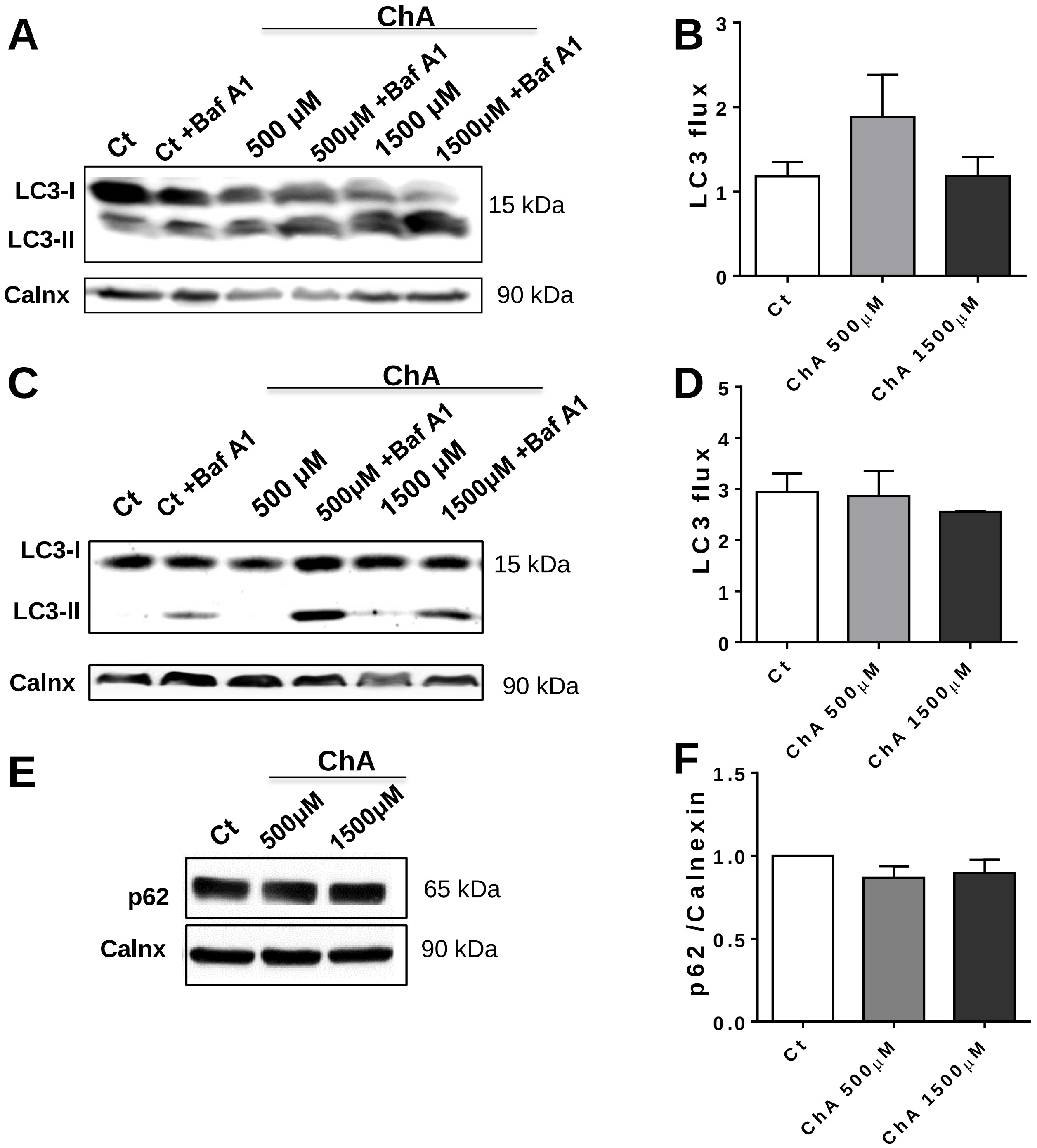


**Supplementary Figure VII: ChA does not alter autopahgy in RAW cells. A.** Immuno-blot of LC-3B in cell lysates of control and ChA-treated cells for 24h in absence or presence Baf A1. **B.** Quantification of the autophagic flux at 24h. **C and D.** Immuno-blot of LC-3B in cell lysates of control and ChA-treated cells for 72 h, without or with Baf A1 treatment. **D.** Quantification of autophagic flux at 72h. **E.** p62 levels in cell lysates of control and ChA-treated cells for 72 h. **F.** Ratio of p62/Calnexin of quantified bands in control and ChA-treated cells. In **B, D**, and **F** the values are mean ± SEM of at least three independent experiments.

#### Supplementary Figure VIII

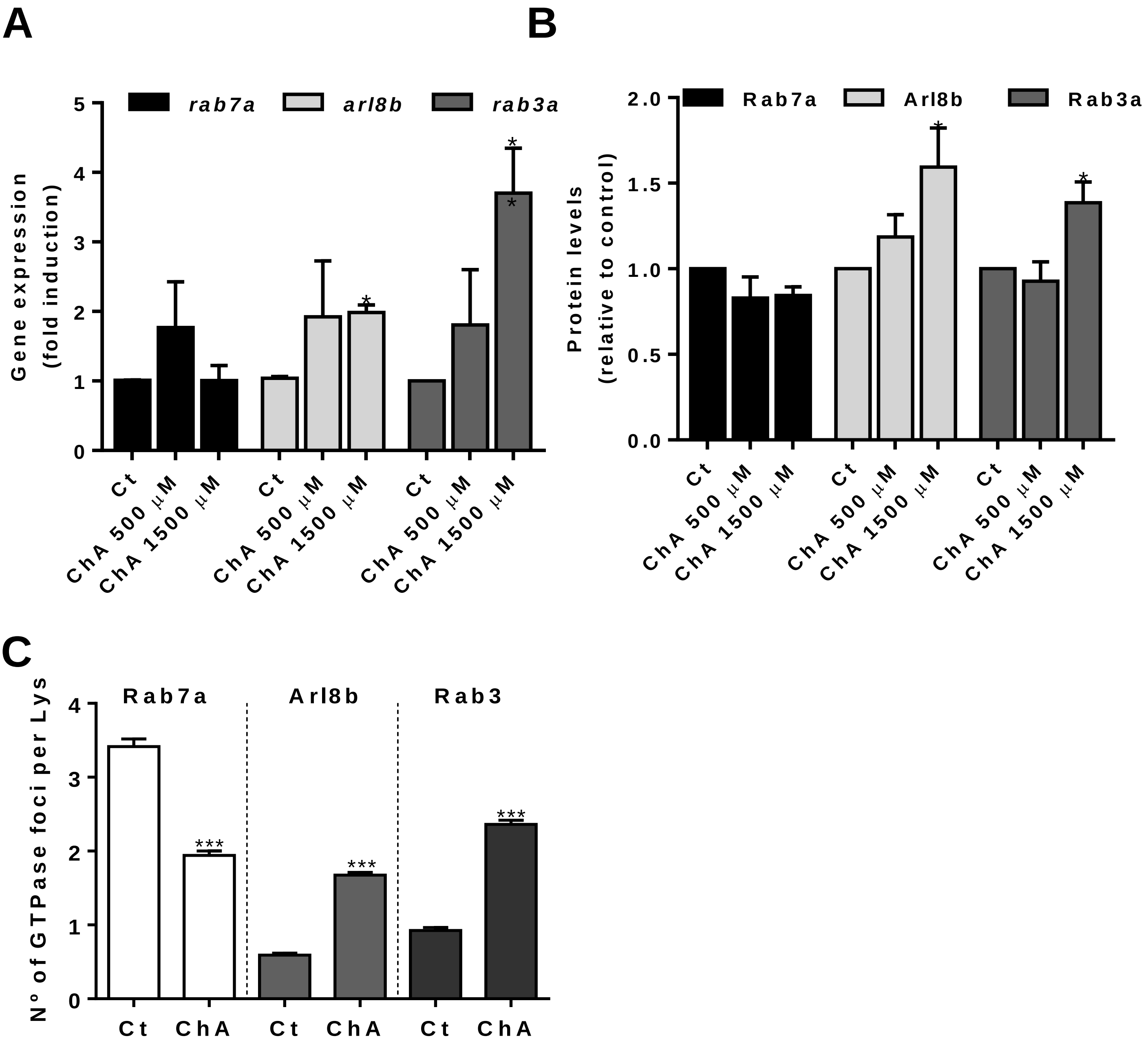


**Supplementary Figure VIII. ChA changes lysosome membrane composition.** RAW cells were incubated for 72 h with POPC or ChA. **A**. *rab7a*, *arl8b* and *rab3a* expression assessed by RT-qPCR. Data were normalized to the endogenous *hprt* and *pgk1* genes. **B.** Quantification by western-blot of the Rab7a/Calnexin, Arl8b/calnexin and Rab3/Calnexin bands obtained from lysates of control and ChA-treated RAW cells. **C.** Quantification, using a developed ImageJ macro, of the number of GTPases foci per LAMP2-vesicle. Rab7a, empty columns. Arl8b, light grey. Rab3, dark grey columns. The values are mean ± SEM of three independent experiments. The *p* values were obtained by one way ANOVA-Kruskal-Wallis test *, p<0.05; ***, p<0.001.

### Supplementary Figure IX

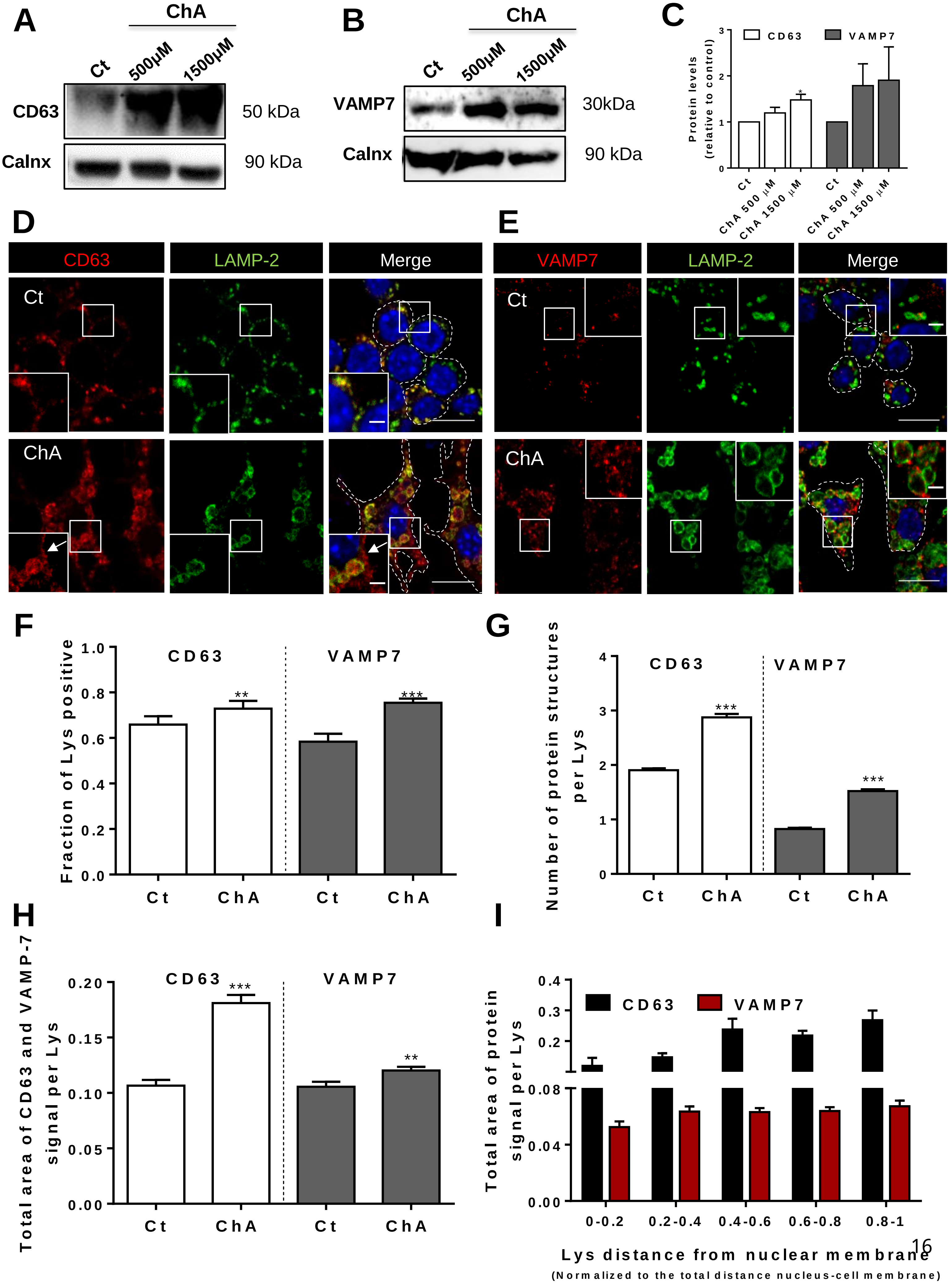

**Supplementary Figure IX. ChA increases the expression of exocytic markers in lysosomal membranes.** RAW cells were incubated with 1500 µM ChA or with POPC (control cells, Ct) for 72 h. Representative western-blots for CD63 (**A**) and VAMP-7 (**B**) from lysates of control and ChA-treated RAW cells. Calnexin was used as loading control. **C.** Quantification by western-blot of the CD63/Calnexin and VAMP-7/calnexin bands obtained from lysates of control and ChA-treated RAW cells. **D.** and **E**. Representative confocal images of control and ChA-treated RAW cells co-immunostained for LAMP-2 (in green) and CD63 (**D**, in red), or VAMP7 (**E,** in red). Nuclei were stained with DAPI (blue). Arrow indicates CD63 staining at the plasma membrane (PM) of ChA-treated cells. The insets are enlargements of the areas outlined with the white boxes. Dashed lines show the edges of RAW cells. Scale bars, 10 µm and 2 µm in the insets. **F.** Quantification of LAMP-2 vesicles positive for CD63 or for VAMP-7. **G.** Quantification of the number of GTPase puncta of CD63 and VAMP7 per lysosome (Lys). **H**. Total area of CD63 or VAMP7 per LAMP-2 vesicles obtained by using a developed ImageJ macro. **I.** Total area of CD63 or VAMP7 per LAMP-2 lysosome (Lys) area as a function of vesicle distance from the nuclear membrane. Values near 1 indicate a close proximity to the PM. Data is the mean ± SEM of three independent experiments. 15 cells were analyzed per condition in each assay (45 cells in total). Statistical significance was assessed by t-test: **, $p<0.01$; ***, $p<0.001$.

#### SUPPLEMENTARY TABLES

**Supplementary Table I –** Information of the important reagents

| Antibody | Vendor or Source | Catalog # | Working concentration | Persistent ID / URL |
|---|---|---|---|---|
| Cholesteryl hemisuccinate | Sigma | 1510-21-0 | NA | https://www.sigmaaldrich.com/catalog/substance/cholesterylhemisuccinate48673151021011?lang=pt&region=PT&gclid=Cj0KCQjw6575BRCQARIsAMp-ksOGrUg8Vq7RmeBIMk1_XKzX2oZgDivSSrmKDUjmyQCiyGbquY8fgMkaAtNGEALw_wcB |
| Cholesterol | Sigma | C8667 | NA | https://www.sigmaaldrich.com/catalog/search?term=57-88-5&interface=CAS%20No.&N=0&mode=partialmax&lang=pt&region=PT&focus=product&gclid=Cj0KCQjw6575BRCQARIsAMp-ksMsw9tz2_OhByux3nXV5sSiQRKD-zuwAe4udLmQCQugNNRMbjZW5sgaArjEEALw_wcB |
| Glutaric anhydride | Sigma | 108-55-4 | NA | https://www.sigmaaldrich.com/catalog/product/aldrich/g3806?lang=pt&region=PT&gclid=Cj0KCQjw6575BRCQARIsAMp-ksMkSslIzhEXEVYaYXn0SwW7v_GbqdP8dufIyzqAae7yAn3QKfYkONQaAk74EALw_wcB |
| 1-palmitoyl-2-oleoyl-glycero-3-phosphocholine (POPC) | Avanti Polar Lipids (Alabaster, AL) | 850457 | NA | https://avantilipids.com/product/850457 |
| TRIzol | ThermoFisher Scientific | 15596026 | NA | https://www.thermofisher.com/order/catalog/product/15596026#/15596026 |
| RNeasy Mini Kit | QIAGEN | 74104 | NA | https://www.qiagen.com/pt/products/discovery-and-translational-research/dna-rna-purification/rna-purification/total-rna/rneasy-mini-kit/?clear=true#orderinginformation |
| DAPI | ThermoFisher Scientific | D1306 | 10 µg/mL | https://www.thermofisher.com/order/catalog/product/D1306#/D1306 |
| BODIPY™ 493/503 | ThermoFisher Scientific | D3922 | diluted 1:100 from a saturated ethanolic solution of Bodipy | https://www.thermofisher.com/order/catalog/product/D3922#/D3922 |
| **DQ-BSA** | ThermoFisher Scientific | D12051 | 50 µg/mL | https://www.thermofisher.com/order/catalog/product/D12051#/D12051 |
| Magic Red | Immunochemistry Technologies | 941 | NA | https://immunochemistry.com/product/magic-red-cathepsin-l-assay-kit/ |
| Bafilomycin | InvivoGen | 88899-55-2 | 62 µg /mL | https://www.invivogen.com/bafilomycin-a1 |
| Dextran conjugated with Alexa Fluor 647 | ThermoFisher Scientific | D22914 | 50 µg/mL | https://www.thermofisher.com/order/catalog/product/D22914#/D22914 |

**Supplementary Table I cont–** Information of the important reagents

| Antibody | Vendor or Source | Catalog # | Working concentration | Persistent ID / URL |
|---|---|---|---|---|
| FITC-dextran | Sigma | FD10S | 250 µg/mL | https://www.sigmaaldrich.com/catalog/product/sigma/fd10s?lang=pt&region=PT |
| Propidium Iodide | Sigma | P4864-10ML | 50 µg/mL | https://www.sigmaaldrich.com/catalog/product/sial/p4864?lang=pt&region=PT&gclid=Cj0KCQjw6575BRCQARIsAMp-ksMITT6B1MLaBuWvQVf5itXTjqBfaeEwsvnZeK10v1_S1OsxeHmtkasaAjKuEALw_wcB |
| FM4-64 | ThermoFisher Scientific | T3166 | 30 µg/mL | https://www.thermofisher.com/order/catalog/product/T13320 |
| NZY first-strand cDNA synthesis kit | NZYtech | mb12501 | NA | https://www.nzytech.com/products-services/molecular-biology/rnacdna/cdna-synthesis/cdna-kits/mb125/ |
| SYBR green master mix | NZYtech | mb22101 | NA | https://www.nzytech.com/products-services/molecular-biology/real-time-pcr/qpcr-master-mixes/mb221/ |
| Azelaic anhydride | Sigma | 246379 | NA | https://www.sigmaaldrich.com/catalog/product/aldrich/246379?lang=pt&region=PT |
| Filipin | Sigma | F9765-25MG | 25 µg/mL | https://www.sigmaaldrich.com/catalog/product/sigma/f9765?lang=pt&region=PT&gclid=Cj0KCQjw6575BRCQARIsAMp-ksOhtq6vo4mxh5HG5mofr4yjUvqD8l-p-S0ohSPOAGzEyhIqU4IbS8QaAg19EALw_wcB |
| BSA-Texas Red | Thermo Fisher Scientific | A23017 | 400 µg/ml | https://www.thermofisher.com/order/catalog/product/A23017 |
| TRITC-transferrin | Thermo Fisher Scientific | T2872 | 200 µg /mL | https://www.thermofisher.com/order/catalog/product/T2872#/T2872 |
| Ionomycin | Sigma | 56092-82-1 | 3.74 mg/mL | https://www.sigmaaldrich.com/catalog/substance/ionomycincalciumsaltfromstreptomycesconglobatus747075609282111?lang=pt&region=PT |

**Supplementary Table II -** List of primary antibodies used for immunoblotting (IB) and immunofluorescent staining (IF)

| Antibody | Vendor or Source | Catalog # | Working concentration | Persistent ID / URL |
|---|---|---|---|---|
| **LAMP- 1** | Developmental Studies Hybridoma Bank | 1-D4B-c | IF: 3.29 μg/ml<br>IB: 0.329 μg/ml | https://dshb.biology.uiowa.edu/1D4B |
| **LAMP- 2** | Developmental Studies Hybridoma Bank | ABL-93-c | IF:2.32 μg/ml<br>IB: 0.232 μg/ml | https://dshb.biology.uiowa.edu/ABL-93 |
| **EEA-1** | Santa Cruz | SC-6415 | IF: 2.00 μg/ml | https://www.scbt.com/p/eea1-antibody-n-19 |
| **Cathepsin D** | Sicgen | AB0043-200 | IF: 15 μg/ml<br>IB: 6 μg/ml | http://www.sicgen.pt/product/cathepsin-d-polyclonal-antibody_1_122 |
| **Rab3** | Sicgen | AB10032-200 | IF: 15 μg/ml<br>IB: 1.5 μg/ml | http://www.sicgen.pt/product/rab3-pan-polyclonal-antibody_1_79 |
| **Rab7** | Abcam | ab50533 | IF: 20 μg/ml | https://www.abcam.com/rab7-antibody-rab7-117-late-endosome-marker-ab50533.html |
| | Sicgen | AB0033-200 | IB: 3 μg/ml | http://www.sicgen.pt/product/rab7a-polyclonal-antibody_1_93 |
| **Arl8b** | made *in house* by YenZym | unknown | IF: 3.3 μg/ml<br>IB: 0.6 μg/ml | NA |
| **CD63** | MBL | D263-3 | IF: 10 μg/ml<br>IB: 2 μg/ml | https://www.mblintl.com/products/d263-3/ |
| **VAMP7** | not applicable (made *in house* by Andrew A. Peden) | unknown | NA | NA |
| **ADRP** | Progen Biotechnik, Heidelberg, DE | GP40 | NA | https://www.progen.com/anti-perilipin-2-n-terminus-aa-1-29-guinea-pig-polyclonal-serum.html |
| **LC3b** | Cell Signaling | 2775S | NA | https://www.cellsignal.com/products/primary-antibodies/lc3b-antibody/2775 |
| **p62/SQSTM1 (2C11)** | Abnova | H00008878-MO | IB: 0.2 μg/ml | http://www.abnova.com/products/products detail.asp?catalog_id=H00008878-M01 |
| **calnexin** | Sicgen | AB0041-200 | IB: 2μg/ml | http://www.sicgen.pt/product/calnexin-polyclonal-antibody_1_30 |

NA- unknown antibody stock concentration according to the manufacter.

**Supplementary Table III -** List of secondary antibodies used immunofluorescent staining

| Antibody | Vendor or Source | Catalog # | Working concentration | Persistent ID / URL |
|---|---|---|---|---|
| Cy™3 AffiniPure Donkey Anti-Rabbit IgG (H+L) | Jackson Immuno Research | 711-165-152 | 1 μg/ml | https://www.jacksonimmuno.com/catalog/products/711-165-152 |
| Donkey anti-Goat IgG (H+L) Cross-Adsorbed Secondary Antibody, Alexa Fluor 488 | ThermoFisher Scientific | A-11055 | 2 μg/ml | https://www.thermofisher.com/antibody/product/Donkey-anti-Goat-IgG-H-L-Cross-Adsorbed-Secondary-Antibody-Polyclonal/A-11055 |
| Cy™3 AffiniPure Donkey Anti-Goat IgG (H+L) | Jackson Immuno Research | 705-165-003 | 2 μg/ml | https://www.jacksonimmuno.com/catalog/products/705-165-003 |
| Cy™3 AffiniPure Donkey Anti-Mouse IgG (H+L) | Jackson Immuno Research | 715-165-150 | 1 μg/ml | https://www.jacksonimmuno.com/catalog/products/715-165-150 |
| Cy™3 AffiniPure Donkey Anti-Rat IgG (H+L) | Jackson Immuno Research | 712-165-153 | 1 μg/ml | https://www.jacksonimmuno.com/catalog/products/712-165-153 |
| Donkey anti-Rat IgG (H+L) Highly Cross-Adsorbed, Alexa Fluor 488 | ThermoFisher Scientific | A-21208 | 2 μg/ml | https://www.thermofisher.com/antibody/product/Donkey-anti-Rat-IgG-H-L-Highly-Cross-Adsorbed-Secondary-Antibody-Polyclonal/A-21208 |
| Cy™5 AffiniPure Donkey Anti-Rat IgG (H+L) | Jackson Immuno Research | 712-175-153 | 1 μg/ml | https://www.jacksonimmuno.com/catalog/products/712-175-153 |
| Donkey anti-guinea pig A488 | Jackson Immuno Research | 706-545-148 | 1 μg/ml | https://www.jacksonimmuno.com/catalog/products/706-545-148 |

**Supplementary Table IV.** Primers sequences for qRT-PCR.

| Gene | Primers sequence |
|---|---|
| *tfeb* | AGGAGCGGCAGAAGAAAGAC; CAGGTCCTTCTGCATCCTCC |
| *lamp1* | ACATCAGCCCAAATGACACA; GGCTAGAGCTGGCATTCATC |
| *arl8b* | CCGCCATGCTGGCGCTCATCTC;<br>AGGTCTCAGCTTCTCCGGGATT |
| *rab3a* | GCATGAATTCATGATGGCTTCCGCCACAGACTCTCGCTAT;<br>ACTCGTCGACCAGCAAGGTCCATTCGCTTTATTG |
| *rab7a* | CCCCAACACTTTCAAAACCC;TGGCCCGGTCATTCTTGTCC |
| *mitf* | GCAAGAGGGAGTCATGCAGT; AGTTGCTGGCGTAGCAAGAT |
| *tfe3* | CCTGAAGGCATCTGTGGATT; TGTAGGTCCAGAAGGGCATC |
| *lipa* | GGCTAGAGCTGGCATTCATC; CTAGAATCTGCCAGCAAGCC |
| *pgk1* | ATGGATGAGGTGGTGAAAGC; CAGTGCTCACATGGCTGACT |
| *lc-3* | TGGTCTACGCCTCCCAAGAA; GTGGGTGTCACATCTCTGCC |

**Supplementary Table V.** Polar lipid concentrations in blood plasma indicated by the lipidomic analysis for each of the three cohorts examined.

| Type of lipid | Control (n=52)* | ACS (n=73)* | CVD2 (n=83)* |
|---|---|---|---|
| Lysophosphatidylcholines | 169.3 ± 53.3 | 104.2 ± 33.9 | 109.1 ± 30.4 |
| Lysophosphatidylethanolamines | 8.6 ± 3.0 | 7.7 ± 3.7 | 6.8 ± 3.4 |
| Phosphatidylcholines | 2042.2 ± 567.8 | 1584.6 ± 426.5 | 1676.9 ± 387.3 |
| Ether-PCs | 82.6 ± 21.9 | 50.2 ± 18.4 | 57.1 ± 18.3 |
| Phosphatidylethanolamines | 34.0 ± 17.8 | 24.8 ± 16.1 | 26.6 ± 16.8 |
| Ether-PEs | 42.5 ± 20.9 | 21.9 ± 13.8 | 30.6 ± 15.0 |
| Phosphatidylinositols | 130.2 ± 38.7 | 92.1 ± 27.9 | 102.5 ± 32.7 |
| Sphingomyelins | 184.8 ± 29.9 | 169.5 ± 33.8 | 168.9 ± 34.5 |
| **Total polar lipids:** | ***2694.2*** | ***2055.0*** | ***2178.5*** |
| Cholesterol | 1279.8 ± 258.1 | 1293.5 ± 426.6 | 1245.4 ± 387.7 |
| **Polar Lipids + Cholesterol:** | ***3974.0*** | ***3348.5*** | ***3423.9*** |
| [Chol] / [Polar Lipids]: | ***0.32*** | ***0.39*** | ***0.36*** |

*Values are in µmoles of lipid per liter of plasma

The polar phospholipids form a monolayer that surrounds the apolar lipid [triacylglycerides (2193,1 µM), diacylglycerides (58,4 µM), and cholesteryl esters (4804 µM)] in lipoproteins of the plasma. We have assumed that most of the cholesterol is also dissolved in this surface monolayer. For simplicity we have considered a homogeneous population of these lipidic particles with a radius of 10 nm (the radius of LDL particle), an average volume of each lipid molecule in the polar lipid monolayer of 1.3 $nm^3$ [4] and its surface area of 3.72 Å.[5] Previous work from our group described the kinetics and thermodynamics of the partitioning of dehydroergosterol (as a cholesterol analogue) between an aqueous phase containing serum albumin and lipidic particles (liposomes and lipoproteins). [6, 7] Assuming that the kinetic rate constants and equilibrium association and dissociation constants obtained in that work are applicable to the partitioning of ChA into the lipidic particles, we have used as a model for calculation of the ChA concentration (for values of the kinetic rate constants see Estronca et al., 2007 [6]; and for calculation of the concentration of lipid associated ChA see Eqn. S6, Supplementary Material, Estronca et al., 2014 [7]). We observed that out of the total 1.5 µmoles of ChA per liter of plasma observed in the ACS cohort, 1.48 µmoles are associated with the lipid monolayer resulting in a concentration of 0.56 M of ChA in the surface lipid layer of the lipidic particles. The remaining 0.02 µmoles of ChA per liter of plasma are expected to be associated with the albumin or dissolved in the aqueous phase of the plasma.
Admittedly, dehydroergosterol is probably less polar than ChA due to the hemiester group in the latter, it may be expected that ChA partition into the polar lipid phase to be lesser extent than dehydroergosterol. However, our earlier work has also shown that a considerably polar amphiphile like a lysophosphatidylethanolamine derivative has a lipid phase/aqueous phase partition coefficient of $\approx 10^5$ [8] while the same value for dehydroergosterol is $\approx 10^6$ . [6] Thus, we conclude that the concentration of ChA in the polar lipid phase should lie between 0.05 and 0.5 M.

#### Supplementary Table VI

Gene set enrichment analysis - Representation of the number of the differentially expressed genes included in the gene set, the pvalue and FDR value estimated considering both up and down significant expression for each gene set.

| | No Genes | pvalue | FDR |
|---|---|---|---|
| mmu04610 Complement and coagulation cascades | 14 | 4,59E-02 | 5,26E-02 |
| mmu00061 Fatty acid biosynthesis | 9 | 9,69E-04 | 2,24E-03 |
| mmu04640 Hematopoietic cell lineage | 23 | 7,49E-04 | 2,01E-03 |
| mmu00650 Butanoate metabolism | 10 | 7,22E-04 | 2,00E-03 |
| mmu04060 Cytokine-cytokine receptor interaction | 62 | 6,63E-04 | 1,91E-03 |
| mmu04514 Cell adhesion molecules (CAMs) | 32 | 6,35E-04 | 1,87E-03 |
| mmu04975 Fat digestion and absorption | 11 | 5,17E-04 | 1,73E-03 |
| **mmu04142 Lysosome** | **99** | **5,08E-04** | **1,73E-03** |
| mmu03320 PPAR signaling pathway | 37 | 4,78E-04 | 1,73E-03 |
| mmu04380 Osteoclast differentiation | 84 | 4,52E-04 | 1,73E-03 |
| mmu04145 Phagosome | 92 | 2,57E-04 | 1,38E-03 |
| mmu01040 Biosynthesis of unsaturated fatty acids | 19 | 2,45E-04 | 1,35E-03 |
| mmu04966 Collecting duct acid secretion | 16 | 1,34E-04 | 9,74E-04 |
| mmu04621 NOD-like receptor signaling pathway | 108 | 7,02E-05 | 8,29E-04 |
| mmu00100 Steroid biosynthesis | 12 | 4,35E-05 | 5,40E-04 |
| mmu04623 Cytosolic DNA-sensing pathway | 35 | 2,46E-05 | 4,84E-04 |
